# Multi-scale deep tensor factorization learns a latent representation of the human epigenome

**DOI:** 10.1101/364976

**Authors:** Jacob Schreiber, Timothy Durham, Jeffrey Bilmes, William Stafford Noble

## Abstract

The human epigenome has been experimentally characterized by measurements of protein binding, chromatin acessibility, methylation, and histone modification in hundreds of cell types. The result is a huge compendium of data, consisting of thousands of measurements for every basepair in the human genome. These data are difficult to make sense of, not only for humans, but also for computational methods that aim to detect genes and other functional elements, predict gene expression, characterize polymorphisms, etc. To address this challenge, we propose a deep neural network tensor factorization method, Avocado, that compresses epigenomic data into a dense, information-rich representation of the human genome. We use data from the Roadmap Epigenomics Consortium to demonstrate that this learned representation of the genome is broadly useful: first, by imputing epigenomic data more accurately than previous methods, and second, by showing that machine learning models that exploit this representation outperform those trained directly on epigenomic data on a variety of genomics tasks. These tasks include predicting gene expression, promoter-enhancer interactions, replication timing, and an element of 3D chromatin architecture. Our findings suggest the broad utility of Avocado’s learned latent representation for computational genomics and epigenomics.

## 1 Introduction

The Human Genome Project, at its completion in 2003, yielded an accurate description of the nucleotide sequence of the human genome but an incomplete picture of how that sequence operates within each cell. Characterizing each basepair of the genome with just two bits of information—its nucleotide identity—yielded many critical insights into genome biology but also left open a host of questions about how this static view of the genome gives rise to a diversity of cell types. Clearly, answering these questions required gathering more data.

In the ensuing 15 years, driven by advances in next-generation sequencing, the research community has developed many assays for characterizing the human epigenome. These include bisulfite sequencing for measuring methylation status, DNAse-seq and ATAC-seq for measuring local chromatin accessibility, ChIP-seq for measuring factor binding and histone modifications, RNA-seq for measuring RNA expression, and Hi-C for measuring the 3D structure of the genome. These assays can quantify variation in important biological phenomena across cell types. Accordingly, large consortia such as ENCODE, Roadmap Epigenomics, IHEC, and GTEx, have run many types of assays in many human cell types and cell lines, yielding thousands of epigenomic measurements for each basepair in the human genome. For example, as of May 1, 2018, the ENCODE project (http://www.encodeproject.org) hosts > 10,000 assays of the human genome.

Although these data have deepened our understanding of genome biology, we are still far from fully understanding it. Gene annotation compendia such as GENCODE are now quite mature, but cell-type-specific annotations of chromatin state remain only partially interpretable [1–3]. Other areas of active research include predictive models of gene expression, promoter-enhancer interactions, polymorphism impact, replication timing and 3D conformation (reviewed in [4]).

One part of the analytical challenge arises from the complexity of the genome and its interactions with other physical entities in the cell, but another part stems from biases and noise in the epigenomic data itself. For example, many such data sets exhibit position-specific biases, reflecting variation in local chromatin architecture or GC bias in the sequencer. Furthermore, no high-throughput assay is perfectly reproducible, and run-to-run differences in the same experiment may reflect either biological variation in the cells being assayed or experimental variance arising from sample preparation or downstream steps in the protocol. Finally, many epigenomic assays are highly redundant with one another, and many cell types are closely related to each other, leading to highly redundant measurements.

To address these challenges, we aim to produce a representation of the human epigenome that is dense and information-rich. Ideally, this representation will reduce redundancy, noise, and bias, so that variance in the representation corresponds to meaningful biological differences rather than technical artifacts. Computationally, this goal can be framed as an embedding task, in which we project the observed collection of thousands of epigenomic measurements per genomic position down to a low-dimensional “latent space.” Our aim is to induce a latent representation of the genome that can be used in place of epignomic measurements as input to machine learning models trained to perform a variety of genomic predictive modeling tasks.

To solve this embedding task we combine two mathematical techniques—tensor factorization and deep neural networks. Epigenomic data sets can be represented as a tensor with three orthogonal axes: genomic position, cell type, and assay type. Tensor factorization is thus a natural framework for distilling this data into an informative latent representation [5]. The deep neural network augments this process with the ability to encode nonlinear relationships among the factors and to capture dependencies along the genomic axis at various scales.

In order to learn a latent representation of human epigenomics, we train our model, which we call “Avocado,” to perform epigenomic imputation. This task involves computationally “filling in” gaps in a tensor of epigenomic data, where gaps correspond to experiments that have not yet been run. Using data from the Roadmap Epigenomics Consortium, we demonstrate that Avocado yields imputed values that are more accurate than those produced by either ChromImpute [6] or PREDICTD [5], as measured by multiple performance measures. Avocado’s imputed data also captures relationships between pairs of histone marks more accurately than these previous approaches. For example, Avocado accurately predicts that activating marks in a promoter region are typically mutually exclusive with repressive marks and are coupled with higher transcription of the associated gene.

Our primary hypothesis is that Avocado’s learned representation will be generally useful in the context of a variety of predictive modeling tasks. The idea that representations can be learned in one setting and then used as input for other settings is similar to that of word2vec [7] and is an example of transfer learning. To test this hypothesis, we consider the tasks of predicting gene expression, promoter-enhancer interaction [8], replication timing, and elements known as “frequently interacting regions” (FIREs), defined on the basis of Hi-C data [9]. For each task, we train a supervised machine learning model on each of seven alternate sets of features—experimentally collected epigenomic measurements for the cell type of interest, the three sets of imputed epigenomic assays for the cell type of interest, the latent representation learned by PREDICTD, the latent representation learned by Avocado, or the experimentally collected epigenomic measurements from all cell types and assays contained within the Roadmap compendium. We include the entirety of the Roadmap compendium as a baseline because, while computationally expensive to train machine learning models on, it contains the full set of information used to learn the Avocado latent representation. In almost every case, we observe that models trained using Avocado’s learned latent representation outperform models trained directly on either the primary or imputed data for the cell type of interest. In those remaining cases, the performance of models trained using Avocado’s learned latent representation is similar to models trained using either the primary or imputed data for the cell type of interest. Notably, the models that utilize the Avocado latent representation outperform those that utilize the PREDICTD latent representation in every cell type for predicting gene expression and FIREs. However, we notice that the use of the full Roadmap compendium proves to be a surprisingly difficult baseline to beat, and that it also consistently outperforms using either the primary or imputed data from a cell type of interest. When models trained using the Avocado latent factors are compared to those trained using the full Roadmap compendium, there are some contexts in which models trained on the Avocado latent factors perform best and some cases where models trained on the Roadmap compendium perform best. These results suggest the broad utility of Avocado’s approach to learning a latent representation of the genome, and that this utility is derived in part from compressing epigenomic assay measurements from all cell types at each genomic position, instead of only a single cell type. Additionally, our results suggest that the process used to learn a latent representation can affect their utility, and that our approach yields a more informative representation than the simpler approach adopted by PREDICTD.

Lastly, we use feature attribution methods to understand the Avocado model. We find that the genomic latent factors encode most of the “peak-like” structure of epigenomic data, while the cell type and assay latent factors serve mostly to sharpen or silence these peaks for a specific track. This observation suggests that the latent representation encodes a rich representation of the functional landscape of the human epigenome.

## 2 Results

### 2.1 Avocado employs multi-scale deep tensor factorization

To produce a latent representation of the genome, we began with the tensor factorization model employed by PREDICTD. In this model, the 3D data tensor is modeled by three 2D matrices of latent factors, corresponding to cell types, assay types, and genomic positions (Fig. 1a). PREDICTD combines these latent factors in a straightforward way by extracting, for each imputed value, the corresponding rows from each of the three latent factors matrices and linearly combining them via generalized dot product operations. Avocado improves upon this approach in two significant ways.

**Figure 1:**
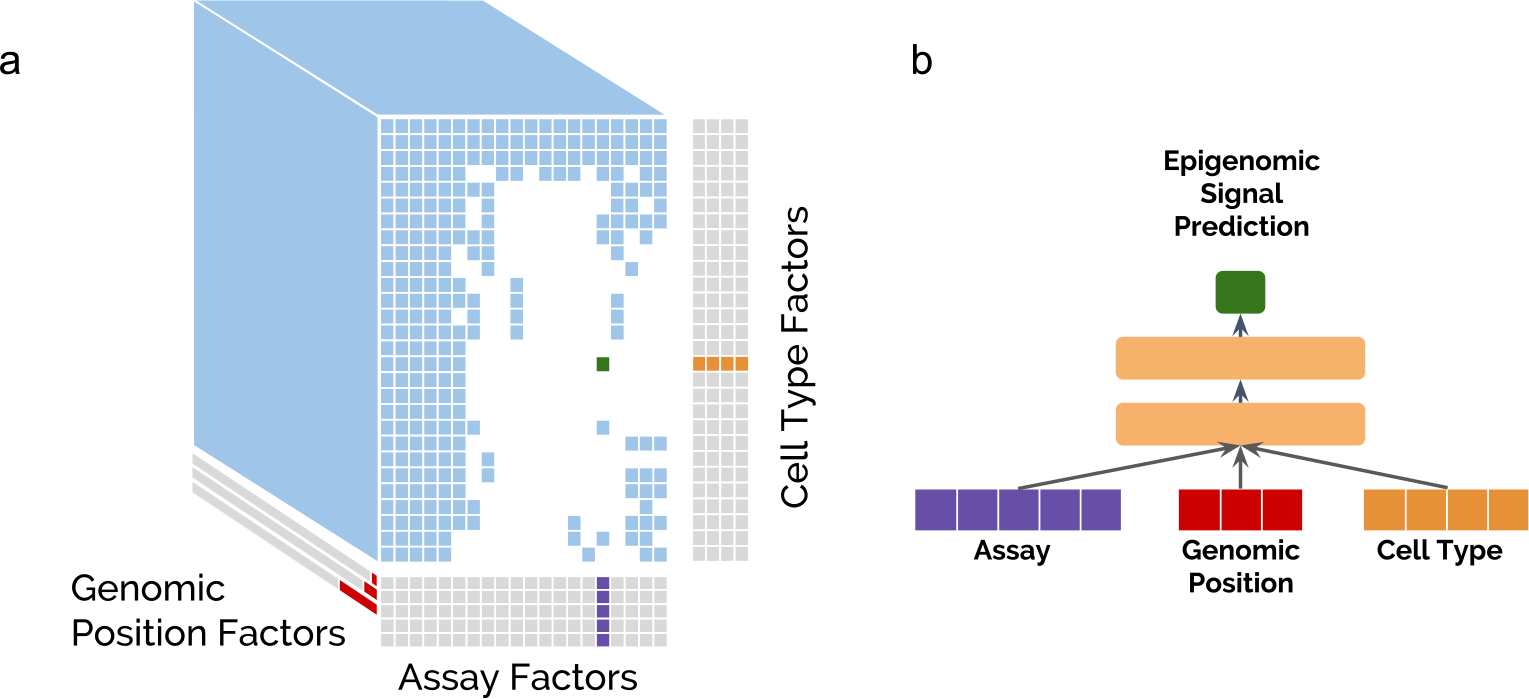
The Avocado deep tensor factorization approach. (a) A collection of epigenomic data can be visualized as a 3D tensor (blue), in which some experiments (white cells) have not yet been performed. Avocado models the tensor along three orthogonal axes, learning latent factors (gray) that represent the cell types (in orange, with 32 factors each), the assay types (in purple, with 256 factors each), and the genomic axis (in red, with 25, 40, and 45 factors at each of the three resolutions). (b) During the training process, the respective slices from these three axes corresponding to the location of the training sample in the tensor are concatenated together and fed into a neural network comprised of two hidden dense layers each with 2,048 neurons to produce the final prediction (in green).

First, Avocado generalizes PREDICTD so that the relationship between the data and the latent factors is nonlinear, by inserting a deep neural network (DNN) into the architecture in place of the generalized dot product operation (Fig. 1b). Note that similar “deep factorization” methods have been proposed previously [10, 11]; however, Avocado differs from these methods in an important way: rather than point-multiplying the three pairs of latent factors and putting the resulting vectors through a DNN, Avocado instead concatenates the three latent factor vectors for direct input to the DNN. This more general approach enables Avocado to embed information about cell types, assay types and genomic positions into latent spaces with different dimensionalities.

**Table 1:** Evaluation of ChromImpute, PREDICTD, and Avocado. Six performance measures are reported, reflecting MSE of different subsets of genomic positions. The best result for each metric is in boldface and corresponds to an adjusted two-sided paired t-test p-value < 0.01 when compared to both other approaches. For MSE-global and MSE-Gene, both PREDICTD and Avocado are bolded because the difference between the two is not statistically significant, i.e., has a p-value > 0.01.

| MSE- | global | 1obs | 1imp | Prom | Gene | Enh |
| --- | --- | --- | --- | --- | --- | --- |
| **ChromImpute** | 0.113 | **0.941** | 1.09 | 0.325 | 0.149 | 0.316 |
| **PREDICTD** | **0.100** | 1.76 | 0.897 | 0.258 | **0.129** | 0.267 |
| **Avocado** | **0.100** | 1.66 | **0.845** | **0.249** | **0.130** | **0.260** |

The concatenation also enables Avocado’s second improvement relative to PREDICTD, namely, that the model adopts a multi-scale view of the genome. Avocado employs three sets of latent factors to represent the genome at different scales: one set of factors are learned for each of the 115,241,319 genomic coordinates at 25 bp resolution, another set are learned at 250 bp resolution, and a final set are learned at 5 kbp resolution. These three length scales represent prior knowledge that important epigenomic phenomena operate at fine scale (e.g., transcription factor binding), at the scale of individual nucleosomes, and at a broader “domain” scale. Furthermore, by learning the genomic representations at multiple scales, Avocado’s genomic latent space can employ far fewer parameters than PREDICTD, requiring only ~3.4 billion parameters instead of ~11.5 billion to model each position along the genome. In total, Avocado requires only ~3.7 percent of the ~92.2 billion parameters employed by PREDICTD’s full ensemble of eight tensor factorization models.

A critical step in developing a model like Avocado involves selecting an appropriate model topology. Avocado’s model (Section 4.2) has seven structural hyperparameters: the number of latent factors representing cell types, assay types, and the three scales of genomic positions, as well as two parameters (number of layers and number of nodes per layer) for the deep neural network. To select these values, we used random search over a grid of hyperparameters, selecting the set that performs best according to the MSE on a validation set when considering the ENCODE Pilot Regions, a selected 1% of the positions in the human genome (Supplementary Note 1). The results of this analysis suggest that, among the seven hyperparameters, the two that control the size of the deep neural network are most important, with Avocado performing best with 2 layers and 2,048 neurons per layer (Supplementary Figure S7). We also found that using “drop-out,” a form of regularization that involves randomly skipping over some model parameters at each training iteration, significantly boosts Avocado’s performance (Supplementary Figure S1).

### 2.2 Avocado imputes epigenomic tracks more accurately than prior methods

We began our analysis of the Avocado latent representation by measuring its ability to impute epigenomic assays, comparing the overall accuracy of Avocado, as measured by mean-squared error (MSE), to that of ChromImpute and PREDICTD. To this end, we evaluated all three methods on 1,014 tracks of epigenomic data from the Roadmap Epigenomics project. Imputations from Avocado and PREDICTD were made using a five-fold cross validation approach, whereas ChromImpute used leave-one-out validation. When considering the full genome, we first evaluated the three approaches using three performance measures originally defined by Durham et al. MSEglobal measures the MSE on the full set of positions; MSE1obs measures the MSE on the top 1% of positions according to ChIP-seq signal; and MSE1imp measures the MSE on the top 1% of positions according to the imputed signal. While Avocado and PREDICTD do equally well according to MSEglobal (unadjusted two-sided paired t-test p-value = 0.451), Avocado outperforms PREDICTD on both MSE1obs (p-value = 9.13e-6) and MSE1imp (p-value = 2.60e-10) (Table 1). This observation is consistent with the observation by Durham et al. that PREDICTD may systematically underpredict signal values, allowing it to score well on regions of low signal but achieving lower accuracy on peaks. Conversely, ChromImpute performs the best on MSE1obs (Avocado/ChromImpute p-value = 2.37e-22, PREDICTD/ChromImpute p-value = 2.85e-12) but the worst on MSE1imp, suggesting that it may over-call peaks. Additionally, Ernst and Kellis proposed six other evaluation performance measures, which show similar trends as the MSE1imp metric (Supplementary Figure S2). We then focused our evaluation on regions of particular biological interest by implementing three more performance measures: MSEProm, MSEGene, and MSEEnh, that measure the MSE of the imputed tracks across all promoter regions, gene bodies, and enhancers, respectively (Table 1). We found that Avocado outperforms the other two methods at MSEProm (Avocado/ChromImpute p-value = 3.98e-32, Avocado/PREDICTD p-value = 8.73e-05) and MSEEnh (Avocado/ChromImpute p-value = 1.72e-30, Avocado/PREDICTD p-value = 1.50e-04), while yielding similar performance to PREDICTD on MSEGene (p-value = 0.875). Taken together, these performance measures suggest that Avocado is able to impute signal well both across the full genome and also at biologically relevant areas (Supplementary Table S1).

All six of the performance measures listed in Table 1 consider each epigenomic assay independently at each genomic position. Empirically, however, many of these assays exhibit predictable pairwise relationships. For example, the activating mark H3K4me3 and the repressive mark H3K27me3 tend not to co-localize within a single promoter region. To measure how well the imputation methods capture such pairwise relationships, we quantitatively evaluated three specific pairwise relationships: negative correlation between H3K4me3 and H3K27me3 in promoter regions, positive correlation between H3K36me3 and RNA-seq in gene bodies, and lack of correlation between H3K4me1 and H3K27me3 in promoter regions. In addition, we considered two pairwise relationships between assays that occur in a promoter and the corresponding gene body: positive correlation between H3K4me3 in promoters and H3K36me3 in the corresponding gene bodies, and the opposite for H3K27me3 and H3K36me3. For each pair of assays, we evaluated how consistent the imputed tracks are with the empirical relationship between the assays (Supplementary Note 2). Across all these evaluations we found that Avocado performed the best at reconstructing the pairwise relationship by between 2.73% and 39.6% when compared to ChromImpute, and between 2.89% and 6.64% when compared to PREDICTD, with PREDICTD typically coming in second and ChromImpute coming in last.

We hypothesized that a primary source of error for all three imputation methods comes from the difficulty in predicting peaks that occur in some cell types but not others. Accordingly, for each assay we segregated genomic positions into those for which a peak never occurs, those in which a peak always occurs (“constitutive peaks”), and those for which a peak occurs in some but not all cell types (“facultative peaks”). Intuitively, we expect an algorithm to be able to predict non-peaks or constitutive peaks more easily than predicting facultative peaks. We test this hypothesis by evaluating the performance of each of the three imputation techniques at genomic positions in chromosome 20 that vary in the number of cell types for which a peak is observed (Section 4.5). This evaluation consists of calculating the MSE, the recall (proportion of true peaks that are imputed), and the precision (proportion of imputed peaks that are true peaks). The recall and precision are calculated by thresholding either the primary or imputed signal at a value of 1.44, corresponding to a signal p-value of 0.01, and evaluating the recovery of MACS2 peak calls. We can determine if a method over- or under-calls peaks based on the balance between precision and recall.

We find that evaluating the three imputation approaches in this manner explains the discrepancy we observed between the MSE1obs and MSE1imp performance measures. Specifically, we find that ChromImpute routinely achieves the highest recall (measured indirectly by MSE1obs) and that Avocado typically achieves the highest precision (measured indirectly by MSE1imp) in regions that are the most variable (Fig. 2). Interestingly, ChromImpute shows a higher recall but lower precision than thresholding the ChIP-seq signal directly in positions that exhibit a peak in many cell types. This observation suggests that ChromImpute may impute wider peaks that, when thresholded, encompass the entirety of the called peak by MACS2. ChromImpute’s high recall and low precision confirm the hypothesis that ChromImpute is over-calling peaks, and specifically that it is likely to predict a peak at a position that is a peak in another cell type. These results also indicate that Avocado and PREDICTD capture different trends in the model, as Avocado typically has higher recall in facultative peaks and PREDICTD has higher recall in constitutive peaks. This finding suggests that one could consider ensembling the imputations from these approaches to yield even more accurate measurements. Overall, Avocado obtains a balance between precision and recall that frequently allows it to achieve the best MSE.

### 2.3 Avocado’s latent representation facilitates a variety of prediction tasks

Having demonstrated that Avocado’s imputed tracks are of high quality, we next investigate the utility of of Avocado’s latent representation as input to tasks for which the model was not explicitly trained for (Fig. 3). This “transfer learning” approach has been used successfully in other domains, such as natural language processing [12] and computer vision [13], and has been more generally described by Pan and Yang [14]. While many techniques can be described as transfer learning, we use the term to refer to training a model for one task and then applying the model (or components thereof) to some other task. Specifically, we hypothesize that Avocado’s latent representation can serve as a replacement for epigenomic data as the input for machine learning models across a variety of genomic prediction tasks. One reason that transfer learning may be beneficial in this case is that many genomic phenomena are associated with epigenomic signals, and so a representation trained to predict these signals is also likely to be associated with these phenomena.

**Figure 2:**
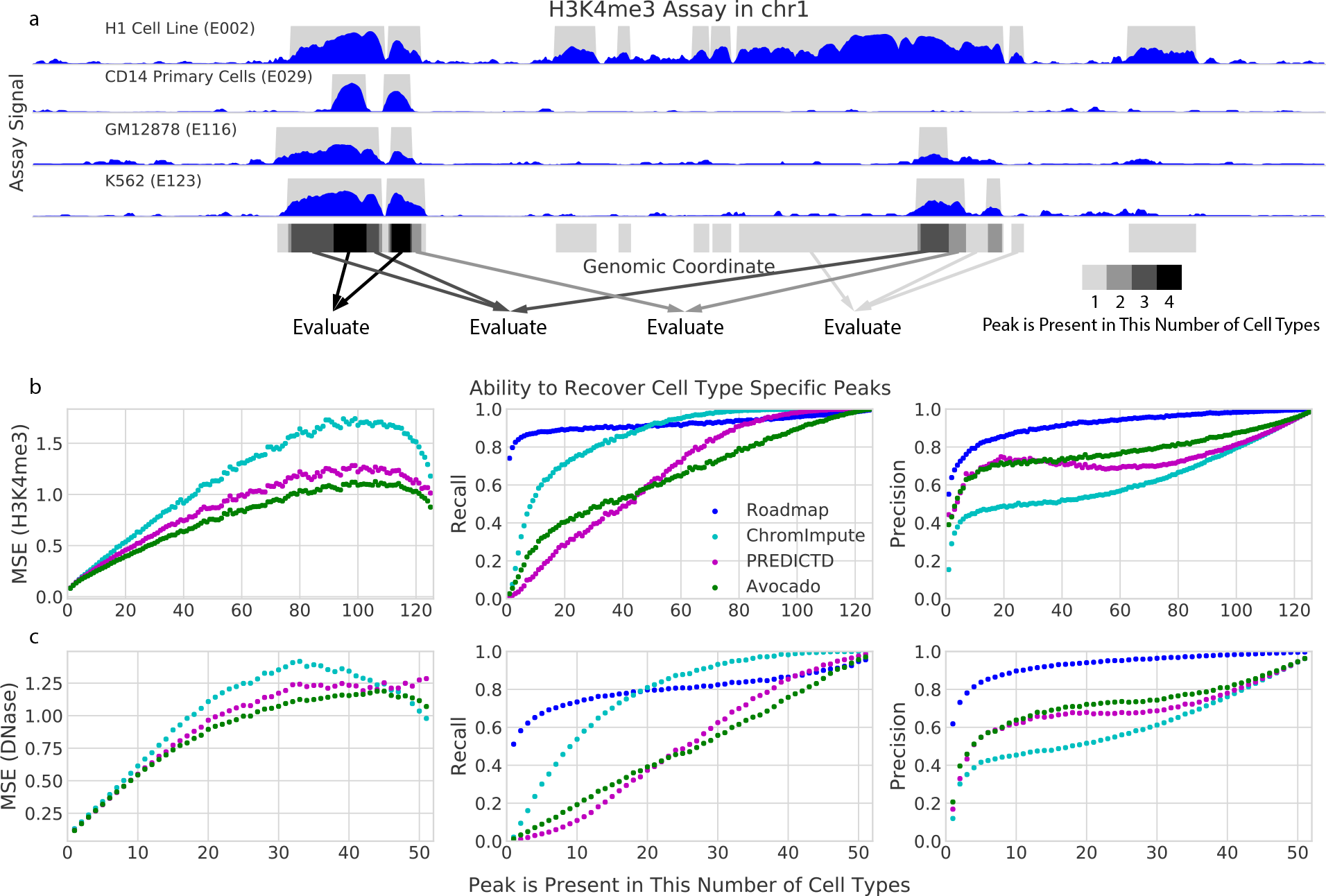
Evaluation of the three imputation approaches at genomic positions that show variation in signal across cell types. (a) A schematic describing how genomic loci are segregated on an example of four cell types. MACS2 peak calls (in gray) are summed over each of the cell type. Genomic loci are then evaluated separately based on the number of cell types in which a peak occurs. (b) Each panel plots a specified performance measure (y-axis) across varying sets of genomic positions (x-axis) for the H3K4me3 assay. For each point, genomic positions are selected based on the number of cell types in which a peak is called at that position, up to a maximum of 127. MSE is calculated between H3K4me3 ChIP-seq signal and the corresponding imputed signal. Precision and recall are computed by thresholding the ChIP-seq signal at 1.44. In the plots, the series labeled “Roadmap” use the observed Roadmap data with peaks called by MACS2. (c) Similar to (b), but using DNAse-seq instead of H3K4me3. All analyses are restricted to chromosome 20.

**Figure 3:**
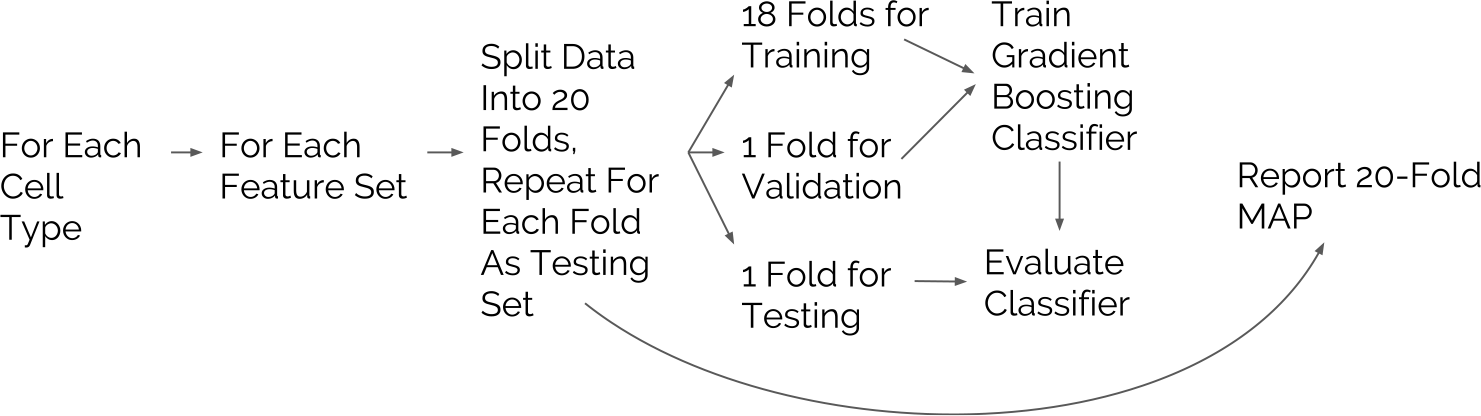
The evaluation procedure for each task. For each cell type and feature set combination, 20-fold cross validation is performed and the MAP across all 20 folds is returned. At each evaluation, a gradient boosted decision tree classifier is trained on 18 of the folds, convergence is monitored based on performance on a 19th fold, and the performance of the resulting model is evaluated on the 20th fold.

We then investigated whether Avocado’s latent representation has implicitly encoded four different types of important biological activity: gene expression, promoter-enhancer interactions, replication timing, and FIREs. These tasks span a diversity of biological phenomena and data sources to ensure that our findings are robust. For each task we train a supervised machine learning model (Section 4.6) using one of seven feature sets: (1) all available ChIP-seq assays for the cell types being considered, (2–4) the set of 24 assays imputed by each of the three methods, (5) the genomic position factors from the single model of PREDICTD’s ensemble that is highlighted in Figure 3 of Durham *et al.* [5], (6) the genomic position factors in Avocado’s latent representation, or (7) the full set of 1,014 ChIP-seq and DNase-seq assays available in the Roadmap compendium (Figure 3). We include the full set of assays from the Roadmap compendium as a baseline feature set because the Avocado latent representation is learned from this full set, allowing us to test our hypothesis that the learned representation preserves cellular variation while removing redundancy and technical noise. Additionally, we include PREDICTD’s learned latent representation to investigate its utility relative to the Avocado latent representation. Lastly, we compare these models to a majority baseline where our prediction for each sample is simply the most prevalent label. We hypothesize that, should the latent representation encode these phenomena well, that the models trained using the latent representation as input will outperform those trained using the other feature sets. Note that the Avocado latent representation is extracted from a model that is trained on the full Roadmap data set. For three of these tasks we use a gradient boosting classifier for this evaluation due to this technique’s widespread success in machine learning competitions [15, 16], with a partial list of top performance on Kaggle competitions available at https://bit.ly/2k7W3Jh.

#### 2.3.1 Gene expression

The composition of histone modifications present in the promoter region of a gene can be predictive of whether that gene is expressed as measured by RNA-seq or CAGE assays. Accordingly, several prior studies have shown that machine learning models can learn associations between these histone marks and gene expression. Because RNA-seq experiments are cheap enough to be performed in any cell type of interest, the typical goal of building a machine learning model is not to replace RNA-seq but to better understand the mechanism behind gene expression. While it may be difficult to explain this mechanism through the interaction of complex latent factors, performing well at this task indicates that complex regulatory information comprised of multiple epigenomic marks is being encoded in the latent factors. Furthermore, a gene expression predictor may be useful in hypothesis generation settings, to assist in prioritizing potential RNA-seq experiments or in investigation of the expression behavior of a small number of genes across many cell types for which epigenomic data has been generated. These studies have approached the problem either as a classification task, where the goal is to predict a thresholded RNA-seq or CAGE-seq signal [17, 18], or a regression task, where the goal is to predict RNA-seq or CAGE-seq signal directly [19].

We approach the prediction of gene expression as a classification task and evaluate the ability of the different feature sets derived from the promoter region of a gene to predict whether or not that gene is expressed. This evaluation is carried out in a 20-fold cross validation setting in each cell type individually, and we report the mean average precision (MAP), which is one technique for calculating the area under a precision-recall curve, across all 20 folds. Genes are considered to be active in a cell if the average RNA-seq value across the gene body is greater than 0.5.

We find that the Avocado latent factors yield the best models in 34 of 47 cell types (Fig. 4a, Supplementary Table S2). In 11 of these 13 cell types, models trained using the Avocado latent factors are only beaten by those trained using the full Roadmap compendium, and in two cell types (E053 and E054; Cortex derived and ganglionic eminence derived neurosphere cultured cells) Avocado is also beaten by models trained using ChromImpute’s imputed epigenomic marks. In no cell type do models trained using the primary data, the typical input for this prediction task, outperform those trained using the Avocado latent representation (unadjusted two-sided paired t-test p-value of 4.62e-153), performing worse by between 0.005 and 0.148 MAP. Additionally, models built using Avocado’s latent representation outperform those built using PREDICTD’s latent representation in every cell type, ranging from an improvement of 0.002 to an improvement of 0.087 (p-value of 3.86e-101). Overall, the models built using the Avocado latent factors perform 0.006 MAP better than those built using the full Roadmap compendium (p-value of 9.75e-21) and only perform 0.001 MAP worse, on average, in those 13 cell types where they perform worse. While this improvement initially appears to be minor, we note that all feature sets yield models that perform extremely well in most cell types, suggesting that there are cell types where gene expression prediction is simple and those in which it is difficult. Accordingly, when focusing on cell types where prediction is more difficult we notice that the difference in performance between the feature sets is more pronounced. Indeed, when we consider the seven cell types where the majority baseline is lowest, we find that those models trained using the Avocado latent factors outperform those trained using the full Roadmap compendium on average by 0.026 MAP and those built using only Roadmap measurements for a specific cell type by 0.107 MAP. These results show that models built using the Avocado latent representation outperform or are comparable to any other feature set considered.

**Figure 4:**
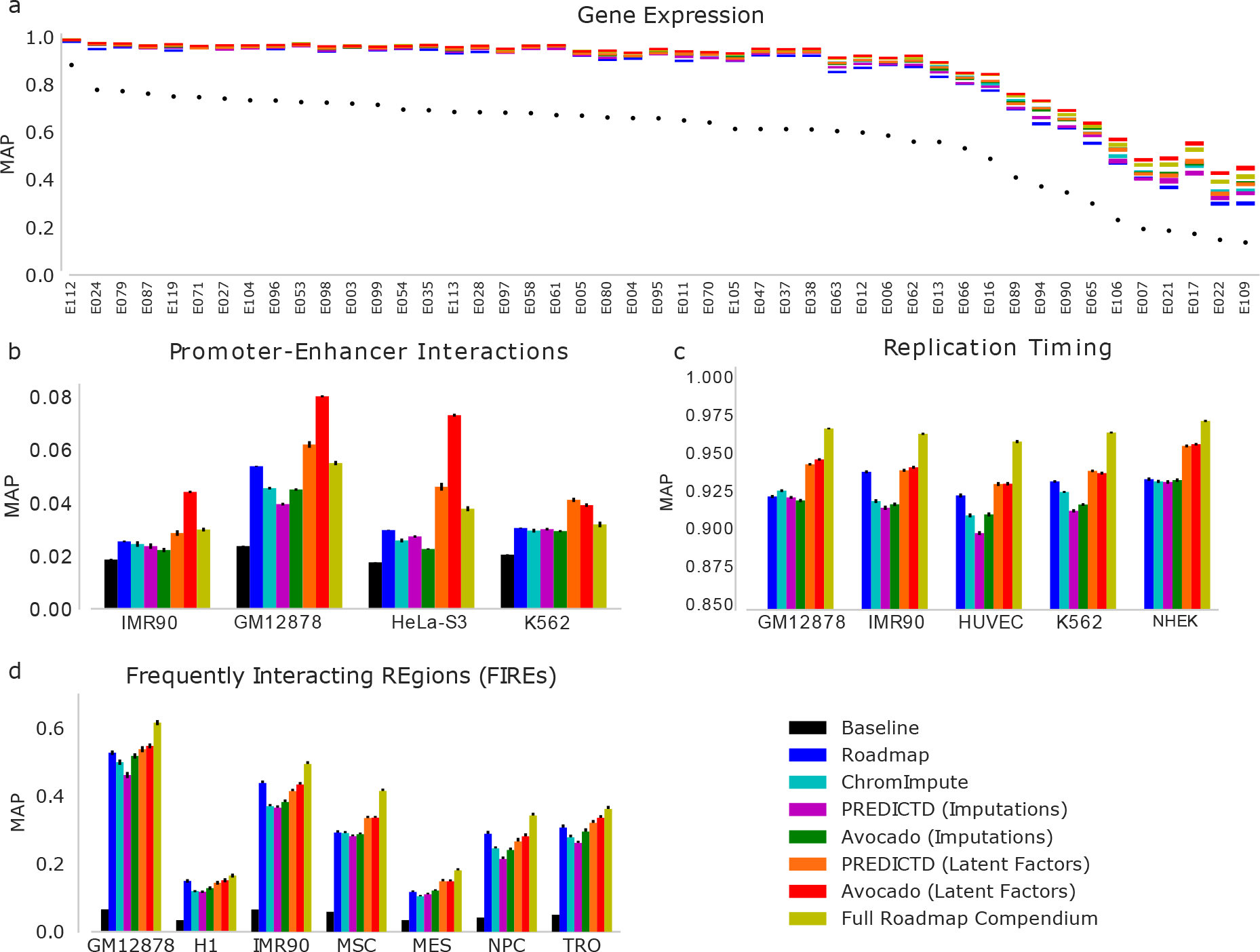
The performance of each feature set when used to for genomic prediction tasks. In each task, a supervised machine learning model is evaluated separately for each cell type using a 20-fold cross-validation strategy, with the mean average precision reported and standard error of the mean shown in the error bars. Each task considers only genomic loci in chromosomes 1 through 22. The tasks are predicting (a) expressed genes, (b) promoter-enhancer interactions, (c) replication timing, and (d) FIREs. In panel (a) the coloring corresponds to the standard error with the mean average precision lying in the middle, whereas in the other panels the mean average precision is shown as the colored bar with standard error shown in black error bars. The statistical significances of differences observed in this figure are assessed in Supplementary Tables S2–S5.

#### 2.3.2 Promoter-enhancer interactions

One of the many ways that gene expression is regulated in human cell lines is through the potentially long-range interactions of promoters with enhancer elements. Physical promoter-enhancer interactions (PEIs) can be experimentally identified by 3C-based methods such as Hi-C or ChIA-PET. However, the resolution of genome-wide 3C methods can be problematic because high resolution contact maps are expensive to acquire. Consequently, predicting PEIs from more widely available and less expensive data types would be immensely valuable. Accordingly, a wide variety of methods for predicting PEIs have been proposed (reviewed by Mora et al. [20]), including those that pair enhancers with promoters using distance along the genome [21], that use correlations between epigenetic signals in the promoter and enhancer regions [22–24], and that use machine learning approaches based on epigenetic features extracted from both the promoter and enhancer regions [8].

We consider the task of predicting physical PEIs as a supervised machine learning problem using features derived from both the promoter and enhancer regions. We employ a set of PEIs that were originally created for training TargetFinder [8], a machine learning model that predicted whether given promoter-enhancer pairs interact with each other using epigenomic measurements derived from both regions. These PEIs correspond to ChIA-PET interactions from each of four cell types (HeLa-S3, IMR90, K562, and GM12878) in chromosomes 1 through 22. We further process this data set to remove a source of bias that has been found since the publication of the original data set [25] (Supplementary Note 3). TargetFinder was not developed to predict interactions in cell types for which contact maps have not been collected, but rather to better understand the connections within existing contact maps. Likewise, we train our classifier to predict PEIs within each cell type, evaluating a regularized logistic regression model in a cross validation setting. For comparison, we use the same collection of real and imputed data types that we used for the gene expression prediction task.

We find that models trained to predict PEIs using the Avocado latent factors perform better than any other feature set that we considered (Fig. 4b) in IMR-90, GM12878, and HeLa-S3. In K562 using the Avocado latent factors is second only to using the PREDICTD latent factors. These improvements in average precision over the full Roadmap compendium range from 0.007 in K562 to 0.035 in HeLa-S3 (p-values ranging from 6.97 × 10^−18^ to 9.45 × 10^−32^, Supplementary Table S3). Interestingly, the PREDICTD latent representation also outperforms the full Roadmap compendium in every cell type (p-value of 2.43 × 10^−22^).

#### 2.3.3 Replication timing

The human genome replicates in an orderly replication timing program, in a process that is associated with gene expression and closely linked to the three dimensional structure of the genome [26, 27]. Patterns of replication timing along the genome can be quantified using experimental assays such as Repli-Seq [28], which can be used to segregate loci into early- and late-replicating regions. Because of the slowly varying nature of replication timing along the genome, we choose to make predictions of early- and late-stage replication at 40 kbp resolution.

Consistent with previous tasks, the Avocado latent representation outperforms both primary and imputed epigenomic data from the cell type of interest (Fig. 4c). However, in contrast to the previous tasks, the Avocado and PREDICTD latent representations perform similarly to each other. While the Avocado latent representation yields models whose improvement over the PREDICTD latent representation is statistically significant (p-value of 0.004, Supplementary Table S4), the effect is small (average precision of 0.9453 vs 0.9442). Further, models that use the full Roadmap compendum yield the best performing models. Taken together, these results suggest that using epigenomic measurements across several cell types can be informative for making predictions even for a single cell type. Additionally, it appears that aggregating these latent spaces to a much coarser resolution (from 25 bp to 40 kbp) may sacrifice valuable information.

#### 2.3.4 Frequently interacting regions

The three-dimensional structure of the genome can be characterized by experimental techniques that identify contacts between pairs of loci in the genome in a high-throughput manner. In particular, the Hi-C assay [29] produces a contact map that encodes the strength of interactions between all pairs of loci in the genome. Within a typical contact map, blocks of increased pairwise contacts called “topologically associating domains” (TADs) segment the genome into large functional units, where the boundaries are enriched for house-keeping genes and certain epigenetic marks such as the CTCF transcription factor [30]. Recently, a related phenomenon, called “frequently interacting regions” (FIREs), has been identified [9]. These regions are enriched for contacts with nearby loci after computationally accounting for many known forms of bias in experimental contact maps. FIREs are typically found within TADs and are hypothesized to be enriched in super-enhancers [9].

Accordingly, we investigate the utility of the Avocado latent representation in predicting FIREs. Our gold standard is derived from Hi-C measurements in seven human cell types at 40 kbp resolution [9]. We frame each task as a binary prediction task, classifying each genomic locus as a FIRE or not. Note that any state-of-the-art predictive model for elements of chromatin architecture would likely include CTCF data, because this mark is highly enriched at structural elements. However, we do not include this factor in our feature set because transcription factors were not included in the Roadmap compendium and thus not used to train the Avocado model. Further, our goal is not to train a state-of-the-art model for predicting FIREs, but to evaluate the relative usefulness of these feature sets.

The results for predicting FIREs are similar to the results from the replication timing task, with models trained using the Avocado latent factors outperforming both those trained using cell type specific epigenomic data (p-value of 6.13 × 10^−8^) and the PREDICTD latent factors (p-value of 2.4 × 10^−4^) (Fig. 4d and Supplementary Table S5). The models trained using the full Roadmap compendium outperform those that use the Avocado latent factors in every cell type except H1 (p-value of 1.85 × 10^−33^). This observation suggests that the inclusion of epigenomic measurements across cell types is important when predicting elements of chromatin architecture, as it was for replication timing, but further suggests that aggregations of these factor values across large genomic loci is not as informative as it was for predicting gene expression or promoter-enhancer interactions.

### 2.4 Avocado’s genomic representation encodes most peaks

We next aim to understand why the Avocado latent representation is such an informative feature set across a diversity of tasks. A well-known downside of neural networks is that they are not as easily interpretable as simpler models due to the larger number of parameters and non-linearities involved in the model. In order to understand these models better, feature attribution methods have recently emerged as a means to understand predictions from complex predictive models. These methods, such as LIME [31], DeepLIFT [32], SHAP [33], and integrated gradients [34], attempt to quantify how important each feature is to a specific prediction by attributing to it a portion of the prediction. A useful property of these attributions is that they sum to the resulting prediction, or the difference between the prediction and some reference value.

We chose to inspect the Avocado model using the integrated gradients method, due to its simplicity, in order to understand the role that the various factors play in making predictions. When we run integrated gradients, the input is the set of concatenated latent factors that would be used to make a prediction at a specific position, and the output is the attribution to each factor for that prediction, specifically, the imputed signal at a genomic position for an assay in a cell type. However, the individual factors are unlikely to correspond directly to a distinct biological phenomena. Conveniently, since the attributions sum to the final prediction (minus a reference value), we can sum these attributions over all factors belonging to each component of the model. This aggregation allows us to divide the imputed signals into the cell type, assay, and the three scales of genome attributions.

**Figure 5:**
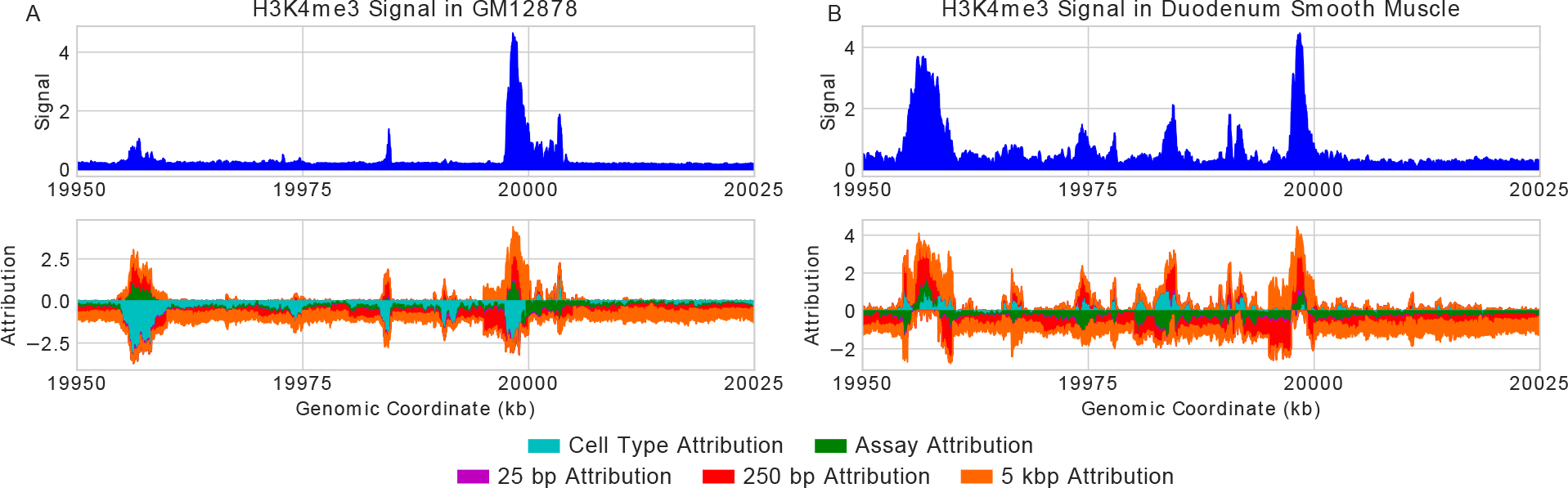
The predicted H3K4me3 signal and corresponding attributions for two cell types in the same region of chromosome 20. (a) The prediction and attributions for GM12878, where a tall peak on the right is paired with two much smaller peaks to the left. Many short regions have a positive genomic attribution but a negative cell type attribution that masks them. (b) The prediction and attributions for deuodenum smooth muscle. A prominent peak is now predicted on the left, corresponding with a swap from a negative cell type attribution to a positive one. The same short regions that previously were masked by the cell type attributions now have positive cell type attributions and exhibit peaks in the imputed signal.

Upon inspection of many genomic loci, most peaks are encoded in the genomic latent factors, while the cell type and assay factors serve primarily to sharpen or silence peaks. An illustrative example of the role each component plays is to consider a pair of nearby regions in chromosome 20 where a H3K4me3 peak with high signal is imputed near a much weaker peak for GM12878 with a very narrow spike between them (Fig 5a) Within the imputed peaks the genome factors predominantly increase the signal, whereas the assay factors appear to increase the signal at the cores of both peaks but dampen the signal on the flanks, effectively sharpening the peaks. Interestingly, the weaker peak appears to have a more prominent signal from the genomic latent factors that is mitigated by a large negative signal from the cell type axis. This indicates to us that this region exhibits a peak in some cell types but is being silenced in GM12878. To confirm that this region engages in a peak in some cell types we looked at the same region in deuodenum smooth muscle cells (E078, Fig 5b) and observed a strong peak (maximum value 3.70 compared to 1.05 in GM12878) that is bolstered by the cell type factors. In addition, there are many smaller peaks that exist in the deuodenum signal that are masked by a negative cell type attribution. This suggests that, while the cell type and assay factors can have positive attributions, they do not fully encode peaks themselves.

We next systematically evaluate the attributions of each component of the model to better understand how Avocado works. Our approach for this analysis is similar to that of analyzing the accuracy of the imputation methods at regions of cellular variability. Specifically, we first segregate positions into bins based on the number of cell types that exhibit a peak at that location, then for each bin we calculate the average attribution in those cell types for which a peak does or does not occur (Supplementary Figure S5). In this manner we can analyze each of the five components of the model in each assay. What we find is that the average cell type attributions are uniformly negative across assays and variability of position when peaks are not present. Additionally, these average attributions typically have a larger magnitude at those variable loci in cell types for which a peak is not present, suggesting that the cell type factors are involved in silencing these peaks in the resulting imputations. The only context in which average cell type attributions are positive are when peaks are present at loci that infrequently exhibit peaks suggesting that the cell type factors may encode infrequent peaks. In contrast, the genomic factors typically have positive values when peaks are present, with negative values correspondingly occuring in infrequent peaks and when peaks are not present. If these rare peaks are a result of technical noise rather than real biology, then this suggests one reason that the genomic factors frequently yield better machine learning models than experimental data.

However, this also suggests that the genomic factors may not be useful at identifying biological phenomena that are indicated by these rare peaks. Interestingly, while the assay attribution values can either be positive or negative, these attributions are higher when peaks are not exhibited rather than when they are. It is unclear why this phenomenon occurs, but it further indicates that the genomic components of the model are a critical driver of Avocado predicting a peak.

## 3 Discussion

Avocado is a multi-scale deep tensor factorization model that learns a latent representation of the human epigenome. We find that, when used as input to machine learning models, Avocado’s latent representation improves performance across a variety of genomics tasks relative to models trained using either experimentally collected epigenomic measurements or the full set of imputed measurements. This representation is more informative than the one learned through the linear factorization approach taken by PREDICTD, suggesting that latent representations can vary in utility and that more work will need to be done to understand them fully. Additionally, in the context of replication timing and FIRE prediction, we found that aggregating both the PREDICTD and the Avocado latent spaces to much lower resolutions by averaging factor values appeared to diminish their utility, suggesting that perhaps these latent spaces are not linearly interpolatable. We have made the Avocado latent representation available for download from https://noble.gs.washington.edu/proj/avocado/.

We hypothesized that a primary reason that this latent representation is so informative is that it distills epigenomic data from all available cell types, rather than representing measurements for only a single cell type. Indeed, feature attribution methods suggest that the genomic latent factors encode information about peaks from all cell types and assays. However, while verifying this hypothesis, we also found that, contrary to common usage, models that exploit the full Roadmap compendium consistently outperform those that use only measurements available in a single cell type. One explanation for this observation is that cellular context can serve as an implicit regularizer for machine learning models, in the sense that the model can learn to discount peaks that appear in exactly one cell type due to experimental noise or technical error.

Although the Avocado latent representation does not outperform using the Roadmap compendium on all tasks, Avocado is much more practical to use. Avocado’s representation consists of only 110 features, whereas the full Roadmap compendium has 1,014 experiments. Accordingly, we observed that models could be trained from Avocado’s learned genomic representation five to ten times faster than those trained using the full Roadmap compendium. This speedup becomes especially important when the input to a machine learning model is not a single genomic window, but multiple adjacent windows of measurements, as is frequently the case when modeling gene expression. For example, if one were to describe a promoter as eight adjacent 250-bp windows spanning ±2 kb from a transcription start site, then the Avocado representation would have only 565 features due to its multi-scale nature, whereas the Roadmap compendium would comprise 8,112 features. We anticipate that the benefits of a low dimensional representation will become even more important once this strategy is applied to even richer data sets, such as the ENCODE compendium, which is composed of >10,000 measurements. This number of measurements would make building machine learning models very difficult.

A natural desire is to inspect the Avocado latent representation in order to better understand the genome. Unfortunately, we found that such inspection was difficult, in part because the latent factors do not individually correspond to meaningful biological phenomena. An avenue for future studies is to better understand these latent factors through methods that aim to connect learned latent spaces to interpretable concepts [35]. Potentially, one might apply a semi-automated genome annotation method like ChromHMM [1] or Segway [36] to the latent representation directly, with the goal of producing a model that can translate the latent representation into a cell type-independent annotation of the genome.

This is not the first time that latent representations have been trained on one task with the goal of being broadly useful for other tasks. For example, word embeddings have been used extensively in the domain of natural language processing. These embeddings can be calculated in a variety of manners, but two popular approaches, GLoVE [37] and word2vec [7], involve learning word representations jointly with a machine learning model that is trained to model natural language. In this respect, these embedding approaches are similar to ours because the Avocado latent representation is learned as a result of a machine learning model being trained to impute epigenomic experiments.

Our approach is not the only approach one could take to reducing the dimensionality of the data. Potentially, one could use a technique like principal component analysis or an autoencoder to project the 1,014 measurements down to 110 dimensions. Alternatively, one might consider using a model similar to DeepSEA [38] or Basset [39] that trains an embedding of the genome jointly with a neural network. However, these types of approaches would not easily allow for transfer learning between cell types, would not allow for the imputation of epigenomic experiments, and would not incorporate information about local genomic context through the use of multiple scales of genomic factors. Furthermore, generalizing an unsupervised embedding approach to make cross-cell type predictions would be difficult, whereas Avocado’s genomic and cell type factors can be combined in a straightforward way to address such tasks.

In this work, we have only explored the Avocado hyperparameter space with respect to the single dataset employed here; thus, generalizing to a new dataset will require repeating this search. Furthermore, in cases where computational efficiency is critical, our results (Supplementary Figure S7) suggest that models with fewer latent factors might perform nearly as well as the full Avocado model. In such settings, it may be sensible to design an objective function for the hyperparameter search that trades off the predictive accuracy of the model versus the model complexity.

We have emphasized the utility of Avocado’s latent genome representation, but the model also solves the primary task on which it is trained—epigenomic imputation—extremely well. In particular, we found that Avocado produced the best imputations when compared with ChromImpute and PREDICTD as measured by five of six performance measures based on MSE for individual tracks, and that these imputed measurements captured pairwise relationships between histone modifications better than either of the other approaches. While investigating why Avocado performed worse than ChromImpute on one of the performance measures, we found that, for all three imputation approaches, much of the empirical error derives from regions where peaks are exhibited in some, but not all, cell types. In the context of identifying which cell types exhibit peaks at these regions of high variability, ChromImpute had the highest recall but the lowest precision, suggesting that it over-calls peaks at a specific region by predicting peaks in more cell types than they actually occur in. In contrast, both Avocado and PREDICTD had lower recall but higher precision, with Avocado frequently managing to balance the two to produce the lowest MSE. Given that these regions are likely the most important for explaining cell type variability, these results suggest that future evaluations of imputation methods should stratify results, as we have done, according to the cell-type specificity of the observed signals. Such investigations might suggest different Avocado hyperparameter settings, focusing on either improved precision or recall, depending upon the end user’s needs.

## 4 Methods

### 4.1 Datasets

The Roadmap ChIP-seq and DNase-seq epigenomic data was downloaded from http://egg2.wustl.edu/roadmap/data/byFileType/signal/consolidated/macs2signal/pval/. Only cell types that had at least five experiments done, and assays that had been run in at least five cell types, were used. These criteria resulted in 1,014 histone modification ChIP-seq tracks spanning 127 cell types and 24 assays. The assays included 23 histone modifications and DNase sensitivity. RNA-seq measurements for 47 cell types were also downloaded for the purpose of downstream analyses, rather than for inclusion in the imputation task. The full set of 24 assays imputed by ChromImpute were downloaded from http://egg2.wustl.edu/roadmap/data/byFileType/signal/consolidatedImputed/, and the full set of 24 assays from PREDICTD were downloaded from the ENCODE portal at https://www.encodeproject.org/.

The specific ChIP-seq measurements downloaded were the −log_10_ p-values. These measurements correspond to the statistical significance of an enrichment at each genomic position, with a low signal value meaning that there is unlikely to be a meaningful enrichment at that position. Tracks that encode statistical significance, such as the −log_10_ p-value of the signal compared to a control track, typically have a higher signal-to-noise ratio than using fold enrichment. Furthermore, to reduce the effect of outliers on the model, Avocado uses the arcsinh-transformed signal

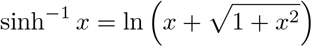

for both training and all evaluations presented here. Other models, such as PREDICTD [5] and Segway [36], also use this transformation, because it sharpens the effect of the shape of the signal while diminishing the effect of large values.

Gene bodies were defined as GENCODE v19 gene elements (https://www.gencodegenes.org/releases/19.html) from chromosomes 1 through 22 that had one of their transcripts annotated as the primary transcript for that gene. This resulted in 16,724 gene bodies per cell type.

Promoter-enhancer interactions were obtained from the public GitHub repository for [8], available at https://github.com/shwhalen/targetfinder/tree/master/paper/targetfinder/combined/output-epw. This data set includes promoter-enhancer interactions as defined by ChIA-PET interactions for four cell lines—GM12878, HeLa-S3, IMR90, and K562. To correct a recently identified bias in this particular benchmark [25], the data set was further processed as described in Supplementary Note 3.

Replication timing data was downloaded from http://www.replicationdomain.org. The resulting tracks encode early- and late-stage timing as continuous values, which are subsequently binarized using a threshold of 0.

FIRE scores were obtained from the supplementary material of [9] for the seven cell lines TRO, H1, NPC, GM12878, MES, IMR90, and MSC. These measurements are composed of binary indicators at 40 kbp resolution, resulting in 72,036 loci for each cell type.

### 4.2 Network topology

Avocado is a deep tensor factorization model, i.e., a tensor factorization model that uses a neural network instead of a scalar product to combine factors into a prediction. The tensor factorization component is comprised of five matrices of latent factors, also known as embedding matrices, that encode the cell type, assay, 25 bp genome, 250 bp genome, and 5 kbp genome factors. These matrices represent each element as a set of latent factors, with 32 factors per cell type, 256 factors per assay, 25 factors per 25-bp genomic position, 40 factors per 250-bp genomic position, and 45 factors per 5-kbp genomic position. For a specific prediction, the factors corresponding to the respective cell type, assay, and genomic position, are concatenated together and fed into a simple feed-forward neural network. This network has two intermediate dense hidden layers that each have 2,048 neurons before the regression output, for a total of three weight matrices to be learned. The network uses the ReLU activation function, ReLU(*x*) = max(0, *x*), on the hidden layers and no activation function on the prediction. The training process jointly optimizes the latent factors in the tensor factorization model and the neural network, rather than switching between optimizing each.

The model was implemented using Keras [40] with the Theano backend [41], and experiments were run using Tesla K40c and GTX 1080 GPUs. For further background on neural network models, we recommend the comprehensive review by J. Schmidhuber [42].

### 4.3 Inputs and outputs

Avocado takes as input the indices corresponding to a genomic position, assay, and cell type, and outputs an imputed data value. The indices for each dimension are a set of sequential values that uniquely represent each of the possibilities for that dimension, e.g., a specific cell type, assay, or genomic position. Any data value in the Roadmap compendium can thus be uniquely represented by a triplet of indices, specifying the cell type, index, and assay.

### 4.4 Training

Avocado is trained using standard neural network optimization techniques. The model was fit using the ADAM optimizer due to its widespread adoption and success across several fields [43]. Avocado’s loss function is the global mean-squared error (MSE). Most training hyperparameters are set to their default values in the Keras toolkit. For the ADAM optimizer, this corresponds to an initial learning rate of 0.01, beta1 of 0.9, beta2 of 0.999, epsilon of 10^−8^, and a decay factor of 1 − 10^−8^. The embedding matrices are initialized with random uniform weights in the range [−0.5, 0.5]. Dense layers are initialized using the “glorot uniform” setting [44]. Using these settings, our experiments show that performance, as measured by MSE, was similar across different model initializations.

Avocado does not fit a single model to the full genome because the genome latent factors could not fit in memory. Instead, training is performed in two steps. First, the model is trained on the selected training tracks but with the genomic positions restricted to those in the ENCODE Pilot Regions [45]. Second, the weights of the cell type factors, assay factors, and neural network parameters are frozen, and the genome factors are trained for each chromosome individually. This training strategy allows the model to fit in memory while also ensuring consistent parameters for the non-genomic aspects of the model across chromosomes, and for the latent factors learned on the genomic axis to be comparable across cell types.

The two steps of training have the same initial hyperparameters for the ADAM optimizer but are run for different numbers of epochs. Each epoch corresponds to a single pass through the genomic axis such that each 25 bp position is seen exactly once, with cell type and assays chosen randomly for each position. This definition of “epoch” ensures that the entire genome is seen the same number of times during training. Training is carried out for 800 epochs on the ENCODE Pilot regions and 200 epochs on each chromosome. No early-stopping criterion is set, because models converge in terms of validation set performance for all chromosomes in fewer than 200 epochs but do not show evidence of over-fitting if given extra time to train.

### 4.5 Evaluation of variable genomic loci

For each assay, we evaluated the performance of Avocado, PREDICTD, and ChromImpute, at genomic positions segregated by the number of cell types in which that genomic locus was called a peak by MACS2. We first calculated the number of cell types that each genomic locus was called a peak by summing together MACS2 narrow peak calls across chromosome 20 and discarded those positions that were never a peak. This resulted in a vector where each genomic locus was represented by the number of cell types in which it was a peak, ranging between 1 and the number of cell types in which that assay was performed. For each value in that range, we calculated the MSE, the recall, and the precision, for each technique. Because precision and recall require binarized inputs, the predictions for each approach were binarized using a threshold on the −log10 p-value of 2, corresponding to the same threshold that Ernst and Kellis used to binarize signals as input for ChromHMM.

### 4.6 Supervised machine learning model training

We performed three tasks that involved training a gradient boosted decision tree model to predict some genomic phenomenon across cell types. In each task, we used a 20-fold cross validation procedure, where the data from a single cell type is split into 20 folds, 19 are used for training and 1 is used for model evaluation. This procedure was performed for each cell type, feature set, and task. These models were trained using XGBoost [46] with a maximum of 5000 estimators, a maximum depth of 6, and an early stopping criterion that stopped training if performance on a held out validation set, one of the 19 folds used for training, did not improve after 20 epochs. No other regularization was used, and the remaining hyperparameters were kept at their default values.

For the task of predicting promoter-enhancer interactions, we used logistic regression as an additional safeguard against the bias issue described in Supplementary Note 3. Rather than perform 20-fold cross-vallidation, we performed 5-fold cross-validation 20 times, shuffling the data set after each cross-validation. We adopted this approach due to the small number of positive samples in each cell type, such that there would be fewer than 10 positive samples in each fold of a 20-fold cross-validation. Additionally, we tuned the regularization strength in the default manner for scikit-learn, which considers 10 regularization strengths evenly spaced logarithmically between 10^−4^ and 10^4^ and choosing the strength that performs best on an internal 3-fold cross-validation on the training set.

We evaluate each model in each task according to the average precision (AP) on the test set, which summarizes a precision-recall curve in a single score. The score is calculated as

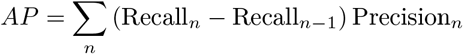

where Recall_*n*_ and Precision_*n*_ are the recall and the precision at the n-th calculated threshold, with one threshold for each data point.

## Data availability

The ChIP-seq and DNase-seq experiments analyzed in this study are currently hosted by the Roadmap Epigenomics Consortium (http://www.roadmapepigenomics.org/) and can be accessed directly at https://egg2.wustl.edu/roadmap/data/byFileType/signal/consolidated/macs2signal/pval/. The data sets generated during this study and the resulting model can be found at https://noble.gs.washington.edu/proj/avocado/. The authors place no restrictions on the download and use of our generated data sets or model.

## Code availability

The Avocado code and documentation are available under the Apache 2.0 License at https://bitbucket.org/noblelab/avocado.

## Acknowledgments

This work was supported in part by an NSF IGERT grant (DGE-1258485) and by NIH awards U54 DK107979 and U41 HG007000.

