## Supplementary Information for "Multi-scale deep tensor factorization learns a latent representation of the human epigenome"

| | Avocado / ChromImpute | Avocado / PREDICTD | PREDICTD / ChromImpute |
|---|---|---|---|
| **MSEglobal** | 1.21e-59 | **4.51e-01** | 2.00e-60 |
| **MSE1imp** | 1.97e-152 | 2.60e-10 | 1.95e-151 |
| **MSE1obs** | 2.37e-22 | 9.13e-06 | 2.85e-12 |
| **GWcorr** | 7.96e-119 | 2.12e-05 | 8.53e-110 |
| **match1** | 1.59e-138 | 1.71e-05 | 3.38e-105 |
| **catch1obs** | 1.04e-154 | 8.45e-50 | 3.79e-90 |
| **catch1imp** | 3.44e-68 | 1.63e-09 | 2.16e-51 |
| **aucobs1** | 5.00e-96 | 3.59e-52 | 4.55e-58 |
| **aucimp1** | 2.60e-25 | 9.22e-04 | 2.29e-18 |
| **MSEProm** | 3.98e-32 | 8.73e-05 | 1.04e-25 |
| **MSEGene** | 1.09e-49 | **8.75e-01** | 7.66e-48 |
| **MSEEnh** | 1.72e-30 | 1.50e-04 | 3.25e-23 |

Table S1: **Statistical significances of imputation performance measures.** Unadjusted p-values from a two-sided paired t-test that compares the average metric value across all 1,014 tracks of data for each pair of imputation methods and performance metric. The two highlighted values are the only two >0.01, indicating that all other comparisons result in statistically significant differences between the two methods.

| | B | R | CI | P(I) | A(I) | P(LF) | A(LF) | FRC |
|---|---|---|---|---|---|---|---|---|
| **Baseline** | — | | | | | | | |
| **Roadmap** | 0.0 | — | | | | | | |
| **ChromImpute** | 0.0 | 5.31e-135 | — | | | | | |
| PREDICTD (I) | 0.0 | 2.48e-27 | 1.34e-99 | — | | | | |
| Avocado (I) | 0.0 | 7.99e-116 | 1.25e-01 | 5.08e-75 | — | | | |
| PREDICTD (LF) | 0.0 | 1.19e-107 | 7.57e-02 | 2.66e-71 | 7.59e-01 | — | | |
| Avocado (LF) | 0.0 | 4.62e-153 | 1.91e-76 | 8.33e-134 | 3.13e-104 | 3.86e-101 | — | |
| FRC | 0.0 | 2.52e-168 | 1.84e-69 | 1.59e-153 | 1.25e-96 | 2.13e-93 | 9.75e-21 | — |

Table S2: **Statistical significances of performance when predicting gene expression.** Unadjusted p-values from a two-sided paired t-test that compares the average precision across all 20 folds from all 47 cell types for a total of 940 measurements. Column names are abbreviated versions of the row names, in the same order. "(I)" stands for "imputations," "(LF)" stands for "latent factors," and "FRC" stands for "full Roadmap compendium." P-values >0.01 are in boldface.

| | B | R | CI | P(I) | A(I) | A(LF) | (LF) | FRC |
|---|---|---|---|---|---|---|---|---|
| **Baseline** | — | | | | | | | |
| **Roadmap** | 2.46e-22 | — | | | | | | |
| **ChromImpute** | 1.15e-23 | 3.29e-10 | — | | | | | |
| **PREDICTD (I)** | 2.82e-32 | 1.66e-08 | **0.0127** | — | | | | |
| **Avocado (I)** | 9.56e-19 | 7.4e-16 | 0.000176 | **0.502** | — | | | |
| **PREDICTD (LF)** | 9.35e-32 | 1.31e-20 | 1.34e-25 | 8.11e-26 | 9.51e-28 | — | | |
| **Avocado (LF)** | 9.45e-32 | 9.54e-26 | 1.53e-26 | 8.33e-27 | 9.32e-27 | 6.97e-18 | — | |
| **FRC** | 1.57e-30 | 2.35e-09 | 2.02e-19 | 1.17e-18 | 2.43e-22 | 1.25e-12 | 1e-24 | — |

Table S3: **Statistical significances of performance when predicting promoter-enhancer interactions.** Unadjusted p-values from a two-sided paired t-test that compares the average precision across all 20 runs from all 4 cell types for a total of 80 measurements. Column names are abbreviated versions of the row names, in the same order. "(I)" stands for "imputations," "(LF)" stands for "latent factors," and "FRC" stands for "full Roadmap compendium." P-values >0.01 are in boldface.

| | B | R | CI | P(I) | A(I) | P(LF) | A(LF) | FRC |
|---|---|---|---|---|---|---|---|---|
| **Baseline** | — | | | | | | | |
| **Roadmap** | 6.91e-143 | — | | | | | | |
| **ChromImpute** | 2.42e-149 | 3.8e-13 | — | | | | | |
| **PREDICTD (I)** | 6.93e-146 | 7.04e-22 | 2.13e-20 | — | | | | |
| **Avocado (I)** | 1.1e-150 | 1.48e-21 | 7.83e-09 | 4.57e-09 | — | | | |
| **PREDICTD (LF)** | 5.37e-154 | 2.35e-22 | 2.73e-62 | 9.98e-75 | 3.77e-83 | — | | |
| **Avocado (LF)** | 5.53e-154 | 2.47e-22 | 4.23e-58 | 6.26e-76 | 1.78e-74 | 0.00406 | — | |
| **FRC** | 6.64e-156 | 5.33e-70 | 3.52e-95 | 5.85e-82 | 1.96e-97 | 1.73e-69 | 2.8e-63 | — |

Table S4: **Statistical significances of performance when predicting replication timing.** Unadjusted p-values from a two-sided paired t-test that compares the average precision across all 20 runs from all 5 cell types for a total of 100 measurements. Column names are abbreviated versions of the row names, in the same order. "(I)" stands for "imputations," "(LF)" stands for "latent factors," and "FRC" stands for "full Roadmap compendium."

| | B | R | CI | P(I) | A(I) | P(LF) | A(LF) | FRC |
|---|---|---|---|---|---|---|---|---|
| **Baseline** | — | | | | | | | |
| **Roadmap** | 3.76e-50 | — | | | | | | |
| **ChromImpute** | 2.89e-47 | 5.04e-21 | — | | | | | |
| **PREDICTD (I)** | 3.17e-48 | 2.80e-29 | 1.18e-08 | — | | | | |
| **Avocado (I)** | 3.79e-48 | 2.17e-12 | 9.62e-06 | 2.15e-17 | — | | | |
| **PREDICTD (LF)** | 7.67e-53 | 4.69e-02 | 6.92e-27 | 1.28e-37 | 1.75e-18 | — | | |
| **Avocado (LF)** | 6.15e-54 | 6.13e-08 | 4.39e-39 | 1.34e-49 | 1.32e-34 | 2.40e-04 | — | |
| **FRC** | 3.54e-56 | 6.72e-41 | 4.26e-62 | 7.37e-61 | 2.07e-55 | 6.94e-39 | 1.85e-33 | — |

Table S5: **Statistical significances of performance when predicting FIREs** Unadjusted p-values from a two-sided paired t-test that compares the average precision across 20 folds from all 7 cell types, for a total of 140 measurements. Column names are abbreviated versions of the row names, in the same order. "(I)" stands for "imputations," "(LF)" stands for "latent factors," and "FRC" stands for "full Roadmap compendium." P-values >0.01 are in boldface.

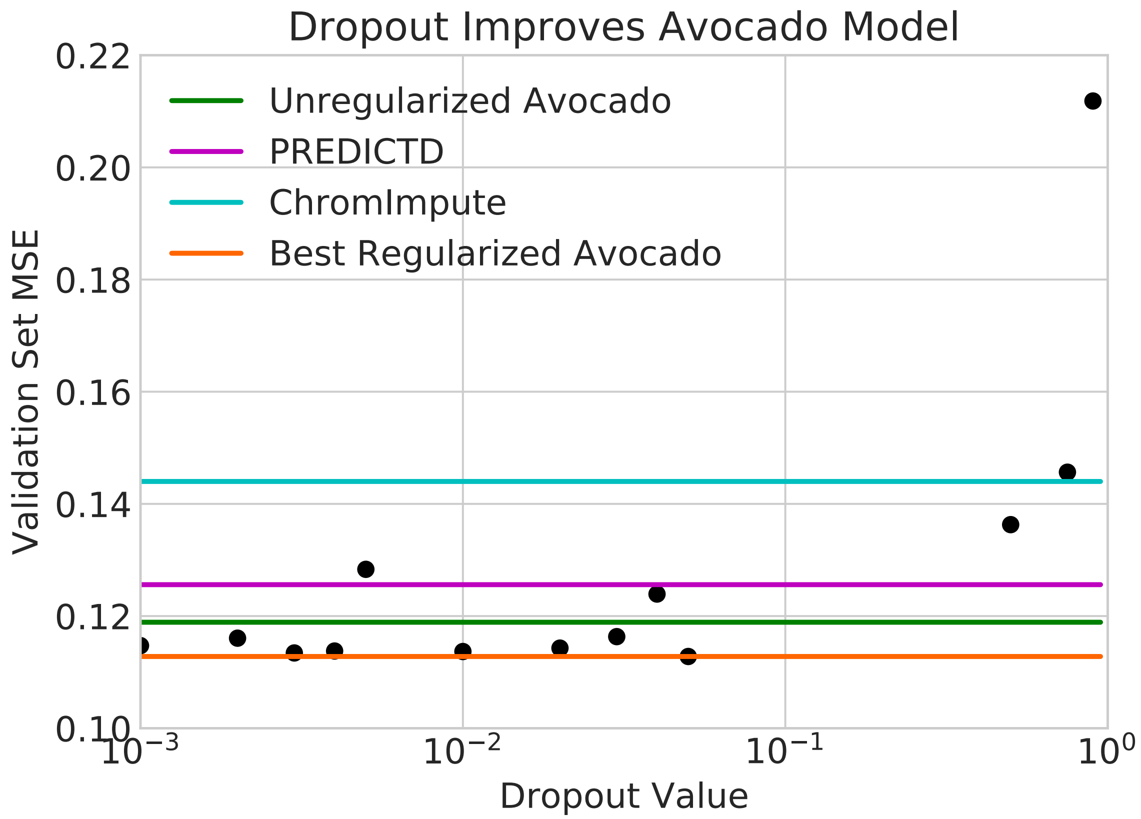


Figure S1: **Dropout improves the validation set performance of Avocado.** Each point corresponds to the performance of an Avocado model trained with a given dropout probability in the two hidden layers. The best performing model (in orange) outperforms not only the unregularized model (in green) but further improves over PREDICTD (in magenta) and ChromImpute (in cyan).

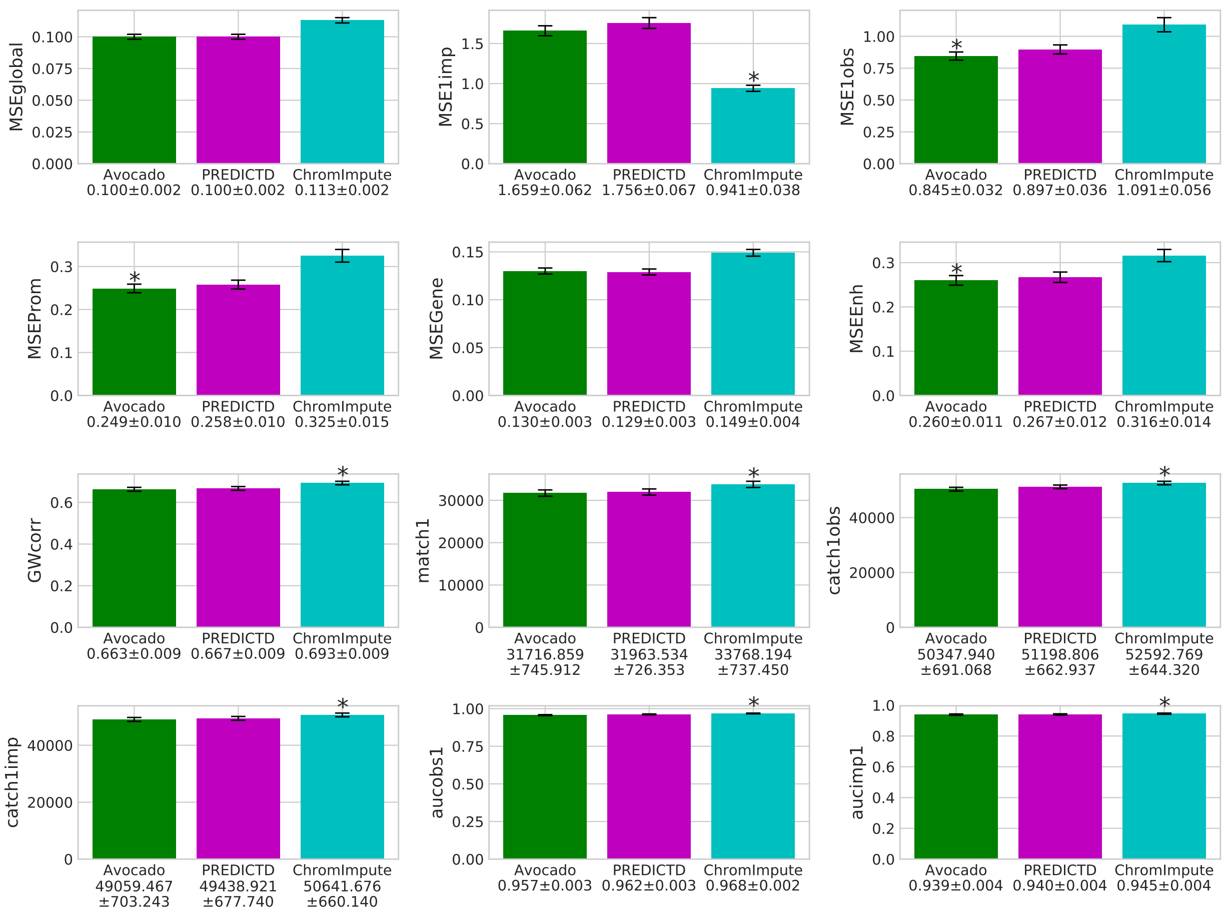


Figure S2: **Twelve performance measures evaluated across the full genome for each imputation approach**. Each panel plots the value of a specified performance measure (y-axis), averaged across all 1,014 tracks. Nine of the performance measures correspond to those proposed by either Durham et al. or Ernst and Kellis. Briefly, MSEglobal is the MSE across the full span of the genome; MSE1imp is the MSE in the top 1% of genomic positions as ranked by the observed signal value; MSE1imp is the MSE in the top 1% of as ranked by the imputed signal value for each approach separately; MSEProm is the MSE of all tracks in promoter regions; MSEGene is the MSE of all tracks in gene bodies; MSEEnh is the MSE of all tracks in enhancers; GWcorr is the Pearson correlation across the full span of the genome; match1 is the number of genomic positions in the top 1% as ranked by observed signal value that are also in the top 1% as ranked by imputed signal value; catch1obs is the number of genomic positions in the top 1% as ranked by observed signal that are in the top 5% of genomic positions as ranked by imputed signal value; catch1imp is as catch1obs but reversed; aucobs1 is the area under the receiver operator characteristics curve (AUROC) when using the imputed signal to recover the top 1% as ranked by observed signal value; and aucimp1 is as aucobs1 but reversed. Error bars display the 95% confidence interval. The best performing approach for each performance measure is denoted with an asterisk above the bar if that result is statistically significant when compared to the next highest performing approach, i.e., p-value $< 0.01$ on a two sided paired t-test, adjusted for the three comparisons.

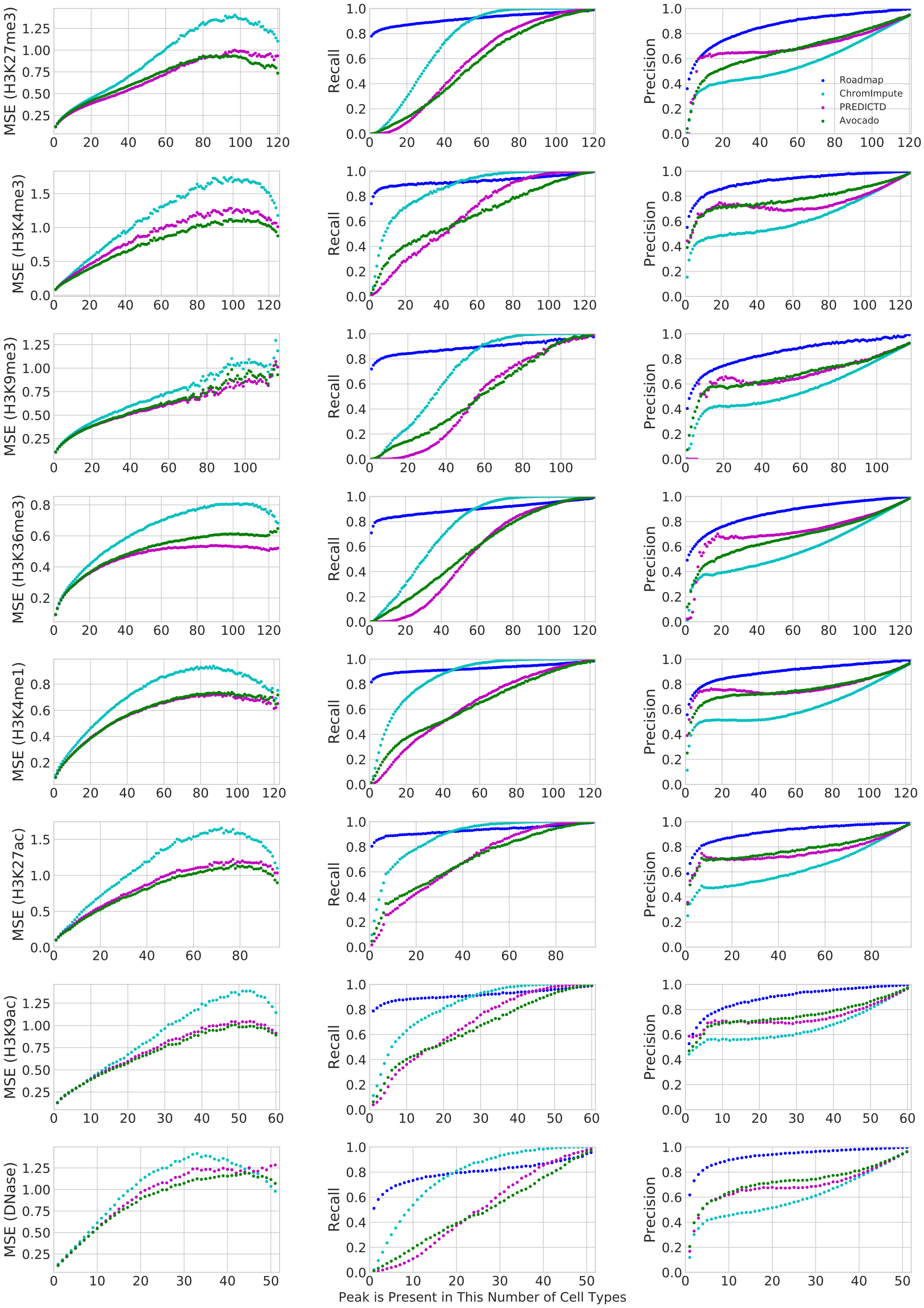


Figure S3: **Ability to recover cell type-specific peaks.** Each panel plots, for a given assay type, the MSE (left column), recall (middle column) or precision (right column) as a function of the number of cell types in which a given peak occurs. Only the 12 assays that have been performed in more than 10 cell types are shown.

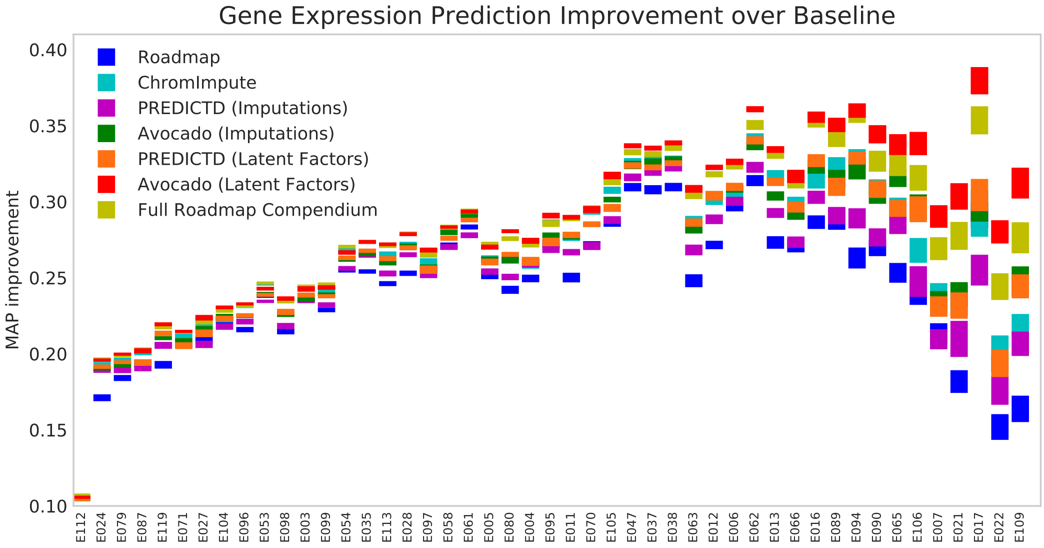


Figure S4: **Relative improvement over a random baseline for each feature set at predicting gene expression.** This plot shows the same values as Figure 4a except that the values for each cell type have the random baseline subtracted out. This view provides a more detailed look at the relative performance of each of the feature sets, even when the performance of all metrics is high.

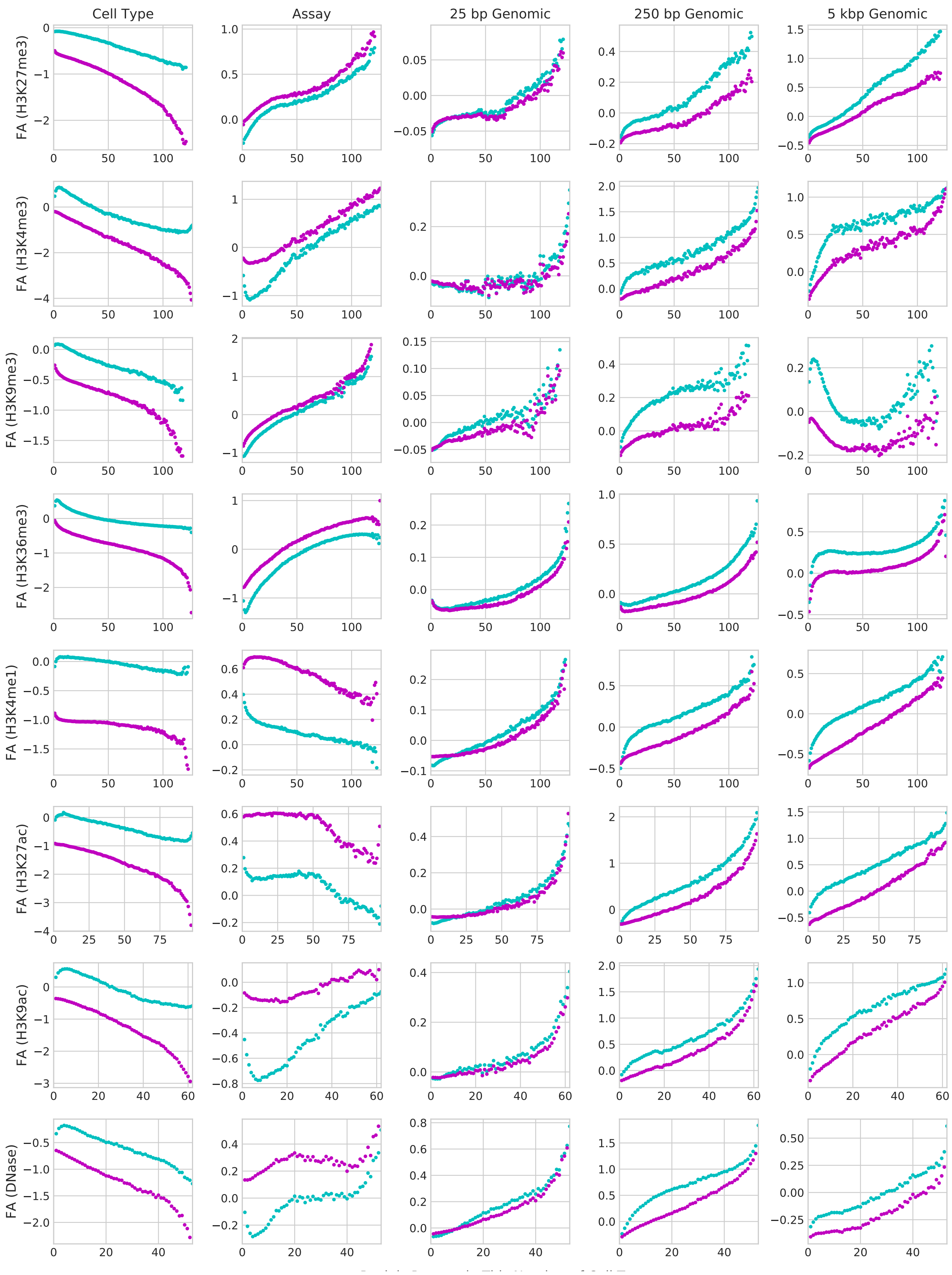


Figure S5: **Feature attribution performed on the Avocado model.** Feature attribution was performed for each position in chromosome 20 across all 1,014 experiments. The results were then aggregated in a manner similar to the analysis of cell-type specific imputations. Instead of calculating the MSE, precision, and recall, instead only the average attribution value is calculated. However, this is done for each of the five model components (the columns). Additionally, the average attribution value is calculated both for those cell types where a peak is exhibited (cyan) and those cell types where a peak is not exhibited (magenta).

### Supplementary Note 1: Hyperparameter Selection

Avocado's model has seven structural hyperparameters: the number of latent factors representing cell types, assay types, and the three scales of genomic positions, as well as two parameters (number of layers and number of nodes per layer) for the deep neural network.

We optimized these hyperparameters via random search. The search considered the following grid of values: cell type factors $\in$ (16, 32, 64, 128, 256), assay factors $\in$ (16, 32, 64, 128, 256), 25 bp resolution genome factors $\in$ (5, 10, 15, 20, 25), 250 bp resolution genome factors $\in$ (10, 20, 30, 40, 50), 5 kbp resolution genome factors $\in$ (15, 30, 45, 60, 75), number of layers in the neural network $\in$ (0, 1, 2, 3, 4), and number of neurons in the neural network $\in$ (128, 256, 512, 1024, 2048). Note that setting the number of layers to 0 corresponds to training a linear regression model on top of the learned factors. These ranges were selected based on experimental results from Durham et al. [1], suggesting that 100 latent factors for each of the three axes performed well. In this grid we trained 1,000 models out of a possible $\sim$61,000. Each model was trained on the ENCODE Pilot Regions, which are comprised of 44 regions of 0.5-2 Mb length that jointly make up approximately 1% of the full genome. The data were split into a training set of 764 tracks, a validation set of 100 tracks, and a test set of 150 tracks. We selected the final set of hyperparameters based on performance on the validation set, as measured by mean-squared error (MSE).

The different hyperparameter settings displayed a wide variance in performance, with most performing better than ChromImpute and many performing better than PREDICTD on the validation set (Supplementary Figure S6). Once the hyperparameters were set, the model was then retrained on both the training and validation sets and tested on the held-out test set. Note that the training, validation, and test sets used here correspond to the same splits used for the PREDICTD approach. The resulting model had a MSE of 0.1130 on the test set, which represents an 18.5% improvement over ChromImpute (MSE 0.1387) and a 4.9% improvement over PREDICTD (MSE 0.1188).

We next investigated the effect that each hyperparameter had on the overall predictive performance of Avocado. To do this, we considered each hyperparameter individually and, for each value that the hyperparameter could take, we plotted the MSE of each model that used that value (Supplementary Figure S7). The clearest trend was that the performance of the model increased as the size of the neural network increased, both in terms of the number of layers and the number of neurons per layer. In contrast, the number of latent factors did not show a clear trend of improvement over any of the three axes.

To attempt to better understand where the allocation of parameters was most beneficial, we considered performance when compared to the total number of parameters in the neural network and when compared to the total number of parameters in the embedding matrices (Supplementary Figure S8). We see that the validation set error decreases steadily with an increase in the number of network parameters until leveling off around $10^7$ parameters. In particular, having no hidden layer, i.e., learning a linear regression on top of the tensor factorization, leads to very poor models. However, adding more than two layers does not yield much gain. When considering the number of parameters at each genomic position in the tensor factorization, we see no similar trend of increased complexity leading to increased performance. We focus on the number of parameters per genomic position rather than the total number of parameters in the model because otherwise the genomic axis would dominate. We can see that having one hidden layer improves model performance; however, we do not see a trend where deeper models are able to better utilize more complex tensor factorization models. Overall, these results suggests that the use of a neural network coupled with the tensor factorization can significantly boost the performance of the model, but that the model is not very sensitive to the complexity of the tensor factorization component.

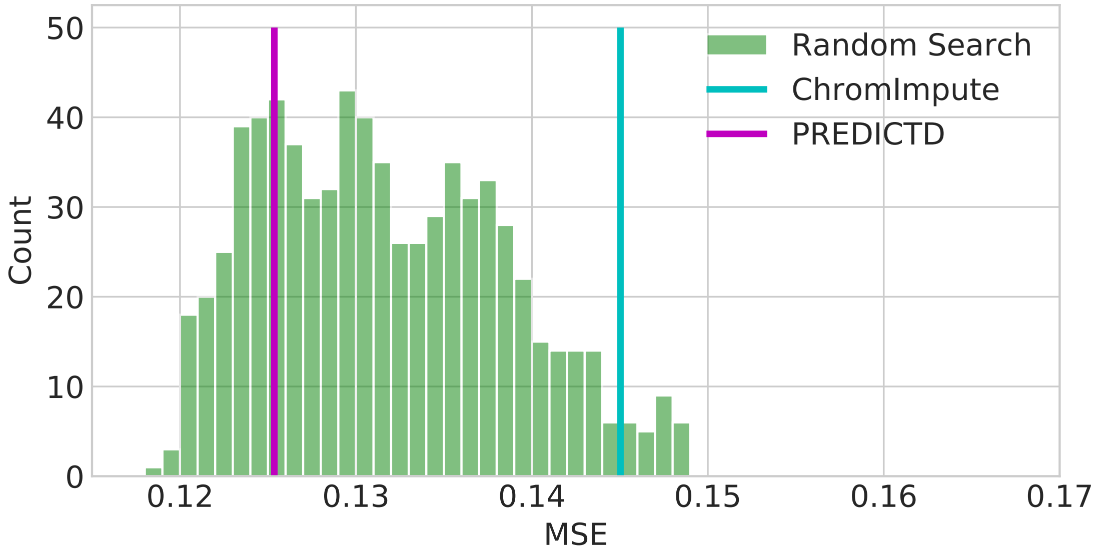


Figure S6: **Random search results on ENCODE pilot regions.** The figure plots a histogram of Avocado validation set MSE values across each hyperparameter setting. For reference, MSE values on the same data set for ChromImpute and PREDICTD are depicted as vertical lines.

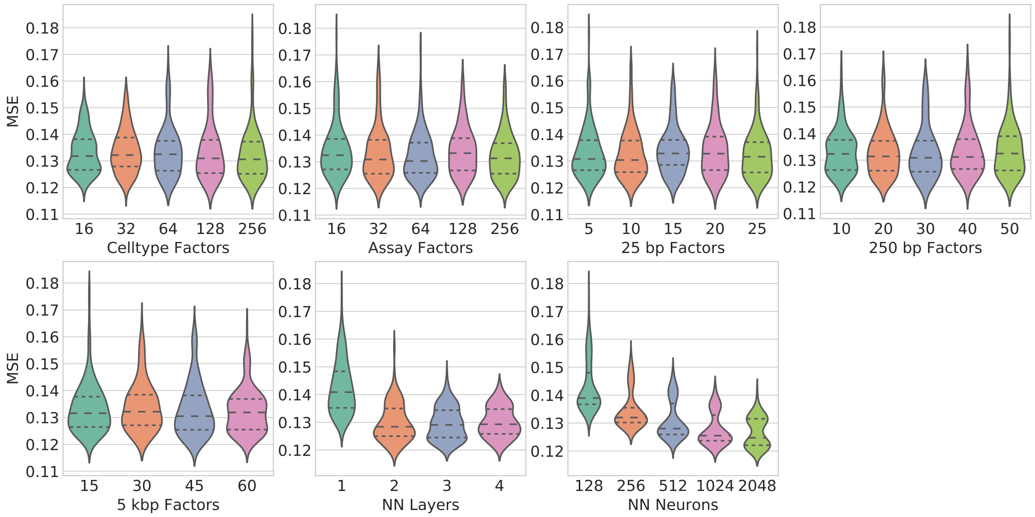


Figure S7: **The performance of the Avocado models learned during random search when stratified by values for each hyperparameter individually.** Each panel shows results for all models that had at least one hidden layer in the neural network. The median is indicated in each violin plot with the longer dashed lines, with the shorter dashed lines indicating the inter-quartile range. The performance seems to be fairly constant across hyperparameter values, except for those hyperparameters related to the neural network. Increasing the number of neurons per layer seemed to increase performance consistently, whereas past two layers the model did not appear to learn significantly more. Models with no hidden layers are not shown, because their performance was uniformly poor.

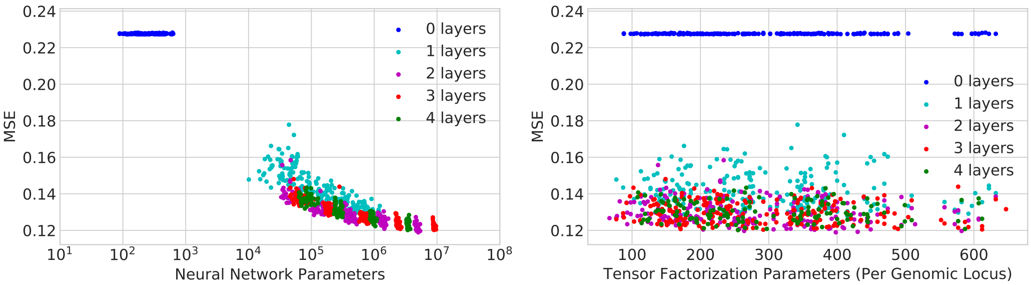


Figure S8: **The number of parameters in each model considered as a part of the random search procedure compared to validation set performance for both the neural network and the tensor factorization aspects.** Left: The trend appears to be that the greater the number of parameters, the better the performance of the model. Models with no hidden layers still have parameters in the form of a linear regression on top of the tensor factorization. The models are colored by the number of layers that they have. Right: The number of parameters in the tensor factorization component at each genomic position. This corresponds to the number of cell type factors plus the number of assay factors plus the number of genomic factors at each resolution. The models are colored by the number of layer in the neural network.

### Supplementary Note 2: Avocado's imputed tracks are consistent with known biology

To better understand the relative behavior of the three imputation methods, we evaluated the imputed measurements for specific histone marks based on their enrichment in functional elements. In particular, H3K4me3 is known to form peaks within transcription start sites (TSSs) and H3K36me3 is known to localize within transcribed genes [2, 3]. We began by extracting the values of H3K4me3 from all TSSs and H3K36me3 from all gene bodies for each cell type. We note that the average H3K4me3 profile across TSSs forms a distinctive bimodal peak (Supplementary Figure S9a). Previously, Ernst and Kellis showed that imputed versions of these histone marks exhibit significantly less variation across cell types than the same signal from ChIP-seq tracks, a trend that is also exhibited by PREDICTD and Avocado [4]. An open question is whether this observed reduced variance corresponds to reduction in noise or reduction in true variation among cell types.

To address this question, we first test whether the observed reduction in variation preserves cellular variation by calculating the rank correlation across cell types between imputed signal and ChIP-seq signal according to the area under each cell types' average mark profile (Supplementary Fig S9a/b). This analysis shows that Avocado preserves the ordering of cell types the best in both H3K4me3 and H3K36me3, while still reducing the variation of the signal. In contrast, while ChromImpute reduces the variation across cell types the most, there is almost no correlation of this measurement between the ChromImpute-imputed H3K36me3 signal and the ChIP-seq measurements. We next test whether cellular variation is maintained by re-implementing the PromRecov and GeneRecov performance measures proposed by Ernst and Kellis that measure how well these two marks localize within their respective regions. All three imputation strategies show similar localization of H3K36me3 in gene bodies (Supplementary Figure S9c), but Avocado shows the highest localization of H3K4me3 in promoter regions in 23 cell types, and a higher localization than ChromImpute in 87 cell types (Supplementary Figure S9d).

To expand on this investigation, we then looked at each techniques' ability to reconstruct relationships among multiple histone marks at the same locus in the genome. We began by looking at the signal values of repressive mark H3K27me3 and the activating mark H3K4me3 in promoter regions, because the two marks tend not to co-localize in differentiated cells (Supp. Figure S10). To quantitatively evaluate this relationship, we calculate the difference between H3K4me3 and H3K27me3 across all 127 cell lines for all promoter regions and calculate the mean absolute error (MAE) between the ChIP-seq signal and the corresponding imputed tracks. This performance measure measures how well the imputation strategies are able to preserve the difference between the two marks. We find that Avocado achieves a lower MAE at reconstructing this relationship than either other method (Supplementary Table S6). We also verified that Avocado does a better job than the other two imputation methods at capturing a lack of correlation between unrelated marks (Supplementary Figure S10), such as the repressive mark H3K27me3 and enhancer-associated mark H3K4me1 (Supplementary Table S6).

We then consider how well the methods can reconstruct the relationship between H3K36me3, a mark typically associated with active gene transcription, and RNA-seq measurements in gene bodies. We restricted our comparison to 47 cell types in which RNA-seq measurements were available from the Roadmap consortium. In this analysis, Avocado captures the relationship the best, and ChromImpute the worst. (Supplementary

| | **ChromImpute** | **PREDICTD** | **Avocado** |
|---|---|---|---|
| **H3K4me4 - H3K27me3** | 0.3566 | 0.3448 | **0.3219** |
| **H3K4me1 - H3K27me3** | 0.3664 | 0.3150 | **0.3059** |
| **H3K36me3 - RNAseq** | 0.2425 | 0.1531 | **0.1464** |
| **H3K4me3 - H3K36me3** | 0.2789 | 0.2887 | **0.2713** |
| **H3K27me3 - H3K36me3** | 0.3163 | 0.2838 | **0.2735** |

Table S6: Evaluation of ChromImpute, PREDICTD, and Avocado at reconstructing relationships between different histone marks across the genome according to the mean absolute error. The best result is in boldface for each comparison.

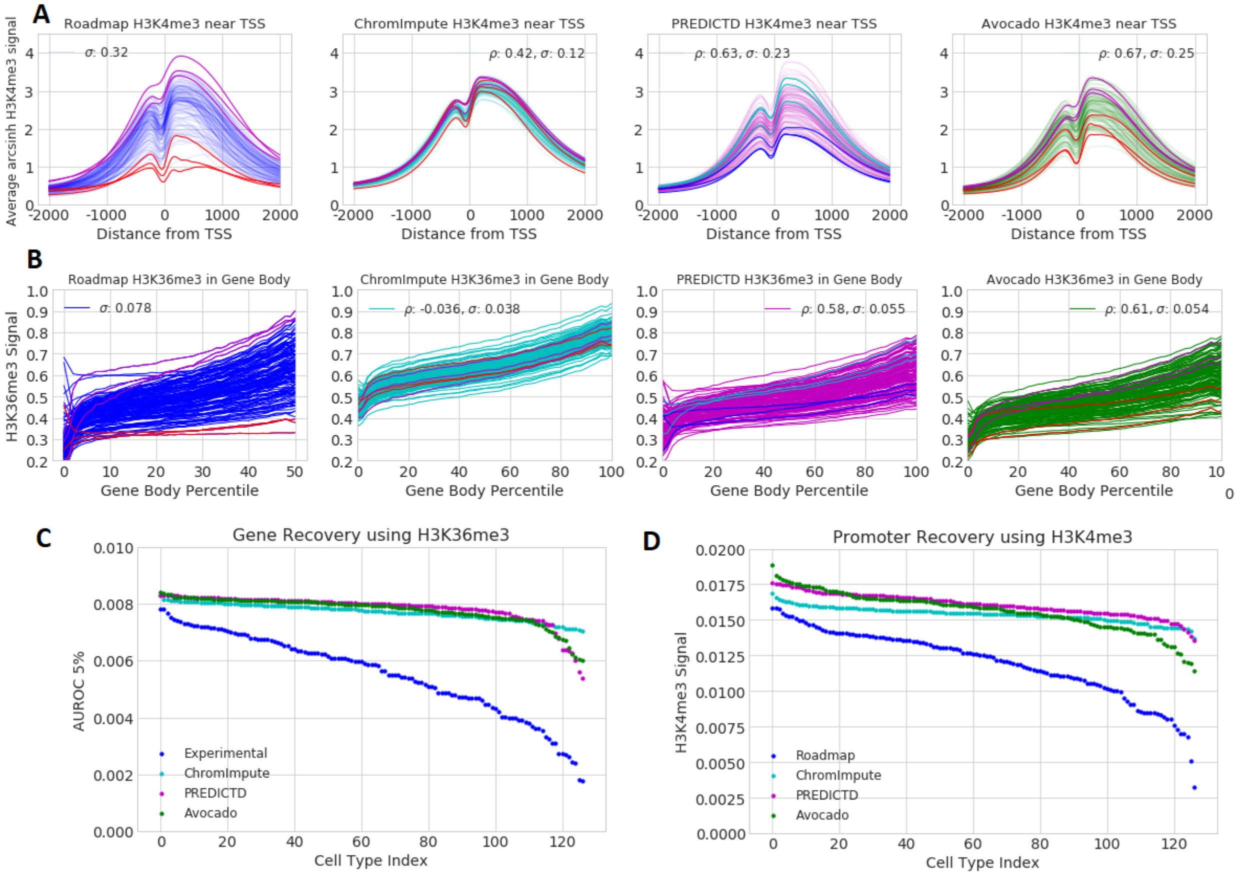


Figure S9: **Aggregate measures of H3K4me3 and H3K36me3 in ChIP-seq experiments and across imputation methods.** (a) H3K4me3 signal in TSSs. Each line displays the average H3K4me3 signal across all TSSs in chromosomes 1-22 for a single cell type after accounting for strand orientation of the gene. The variance of the signal across all cell types at each position is calculated and then averaged ($\sigma$). The area under each line is used to define a ranking, and the spearman correlation ($\rho$) is calculated between each of the three imputation approaches and the ChIP-seq data. To show how the imputation methods alter ordering, the three cell lines with the highest and lowest ChIP-seq signal using this method are colored magenta and red, respectively. In the PREDICTD panels the three with the highest signal are colored cyan and the three with the lowest signal are colored blue. (b) H3K36me3 signal in gene bodies, taking into account the strand orientation of the gene. Measurements are calculated in the same manner as H3K4me3. (c) The GeneRecov performance measure for each cell type. This performance measure quantifies how well H3K36me3 localizes in gene bodies across cell types. It is the area under the ROC curve at 5% FPR when using H3K36me3 to predict gene bodies across chromosomes 1 through 22. (d) The PromRecov performance measure for each cell type. This performance measure quantifies how well H3K4me3 localizes in promoter regions across cell types. It is the area under the ROC curve at 5% FPR when using H3K4me3 to predict promoters across chromosomes 1 through 22.

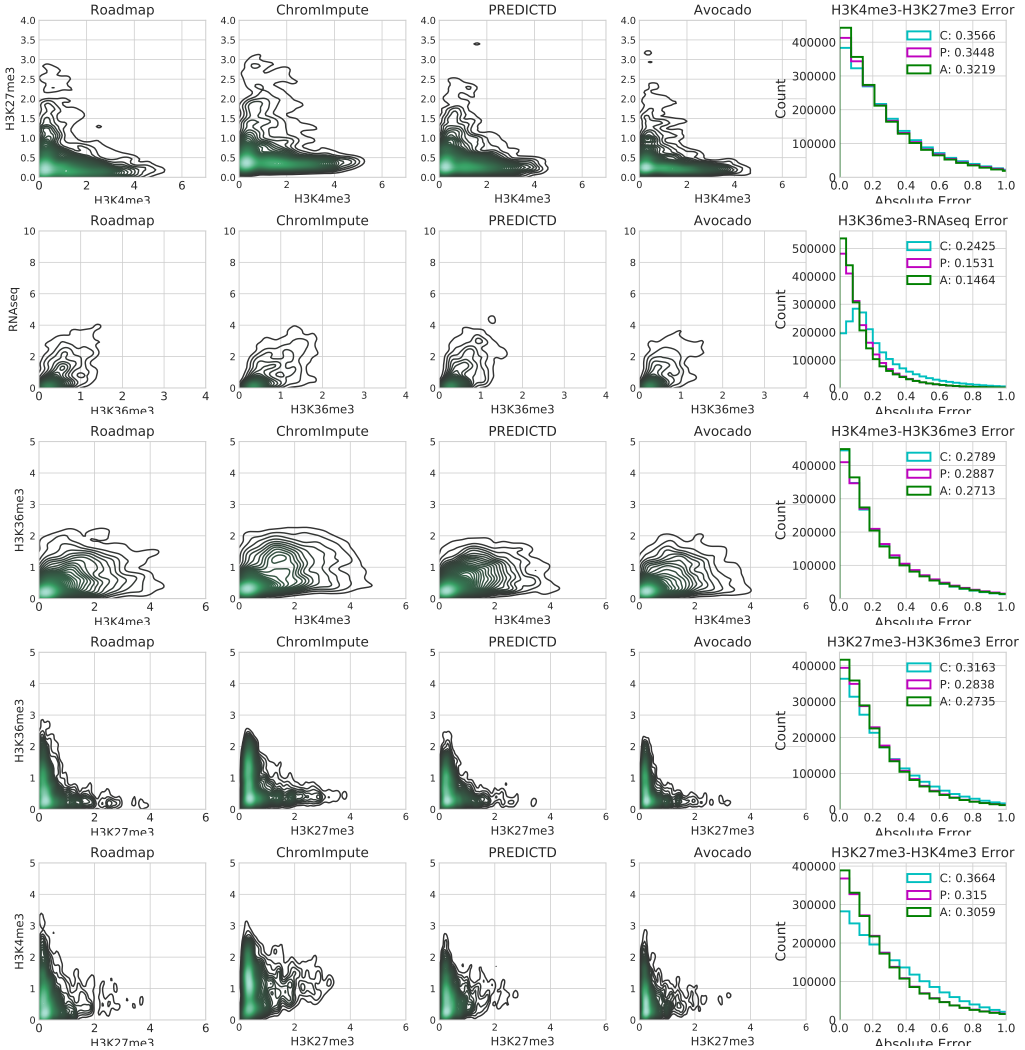


Figure S10: **The relationships between pairs of histone modifications.** These panels show, going from left to right, the signal values in the Roadmap compendium, the imputed signal values from ChromImpute, imputed signal values from PREDICTD, the imputed signal values from Avocado, and the distribution of the absolute error in reconstructing the relationship. In the rightmost panels the legend denotes ChromImpute as C, PREDICTD as P, and Avocado as A. Because each plot contains over 2 million samples, the contour plots are generated on a randomly selected one thousandth of the data, though the error histogram is generated from the full set of samples.

Figure S10).

We then considered relationships across both histone marks and genomic loci, focusing on the relationship between marks in the promoter and the gene body. Specifically, we consider the relationship between H3K4me3 in the promoter region with H3K36me3 in the gene body, because an enrichment of the activating mark should lead to higher levels of the transcription-associated mark. Likewise, we would expect that an enrichment in H3K27me3 in the promoter region should lead to a depletion of H3K36me3 in the gene body. A priori, we expect that ChromImpute and Avocado would do particularly well at reconstructing these interactions because they both take as input information from many nearby genomic loci, whereas PREDICTD treats each genomic position independently. However, we find that while PREDICTD does the worst at reconstructing the relationship between H3K4me3 and H3K36me3, ChromImpute performs much worse at connecting H3K27me3 and H3k36me3 (Supplementary Table S6). Interestingly, despite ChromImpute having an overall negative correlation between H3K27me3 and H3K36me3, as ChromImpute's imputed value of H3K27me3 increases so too does the minimum value of H3K36me3 (Supplementary Figure S10). This trend exists to a much lesser extent in the Avocado model, but is not supported by the ChIP-seq signal.

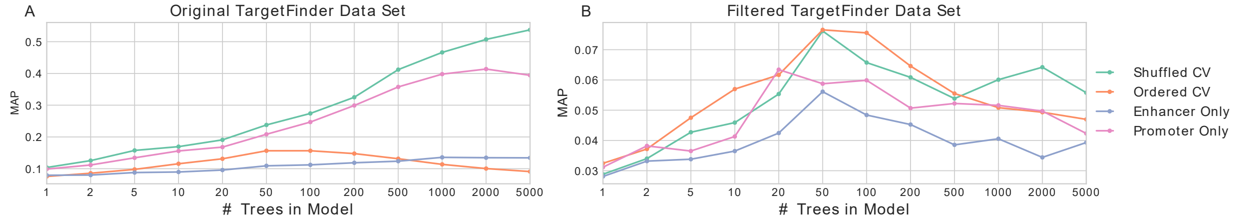


Figure S11: **Model performance on the original and filtered TargetFinder data sets.** (A) The performance of gradient boosting classifiers on the TargetFinder data set split by randomly assigning interactions to folds (cyan) or ordering interactions by genomic coordinate and then splitting into consecutive blocks (orange). Further, when randomly assigning interactions to folds, the performance is shown when using only features from the enhancer (blue) and when using features only from the promoter (pink). (B) Similar to (A), but on the new filtered data set.

### Supplementary Note 3: Promoter-enhancer interaction data set

The set of promoter-enhancer interactions used to evaluate the TargetFinder model [5] has been recently shown to contain biases related to the pairwise nature of the task [6]. The bias arises for two reasons. First, the data set includes features derived from the window between the promoter and the enhancer, and these features are highly correlated between examples whose windows overlap. This correlation leads to a leakage of information when regions of the genome are in windows of examples in both the training and the test set. Fortunately, this issue can be easily corrected by simply removing the problematic features. The second issue is that when the data set was constructed, an equal number of positive and negative interactions were sampled at each genomic distance. Consequently, many promoters occur repeatedly and only in the context of a negative interaction. When promoter-enhancer pairs are randomly assigned to both the training and test sets, as is the case with the TargetFinder model, then a sufficiently complicated model can simply memorize these repeated promoters as never interacting. These issues are described more thoroughly by Xi and Beer [6].

To construct a data set without these biases, we choose the simple approach of filtering out interactions such that each promoter occurs only once in each cell type. While we do not also enforce that enhancers can only occur once, we greedily select pairs where the enhancer has not yet been part of an example. This approach yields a data set with 27,048 interactions across all four cell types in chromosomes 1 through 22, where each interaction corresponds to a unique promoter in its cell type. Among these interactions, nearly all (26,707) have unique enhancers as well; 158 enhancers are seen twice, 7 are seen 3 times, and 1 is seen four times. After this filtering step, IMR90 has 4,702 pairs, of which 82 are positive interactions; GM12878 has 7,881 pairs, of which 181 are positive interactions; HeLa-S3 has 7,060 pairs, of which 121 are positive interactions; and K562 has 7,405 pairs, of which 145 are positive interactions.

We verify that the source of bias has been removed using the same techniques used by Xi and Beer [6]. First, we plot the performance of gradient boosting models with an increasing number of trees evaluated using five-fold cross-validation with examples randomly assigned to folds. We observe a steadily increasing performance on the original data set, similar to that reported by Whalen et al. [5], but not on the new data set (Supplementary Figure S11, orange). Next, we sort examples based on their genomic coordinates and assigned samples to folds based on this ordering. We observe the same diminished performance on the original data set in comparison to random splitting that was observed by Xi and Beer (blue), but similar performance on the new data set compared to random splitting. Lastly, to confirm that this issue is related to memorizing which promoters never interact, we train models using features from only the promoter (green) or the enhancer (brown). We observe similar performance in the original data set when using all features or only using features derived from the promoter region. However, we do not observe this trend in the new data set. Taken together, these results confirm that the new data set does not exhibit the same issue as the original data set used by Whalen et al [5].
